# Identification of RACK1A as a component of the auxin-ethylene crosstalk regulating apical hook development in *Arabidopsis thaliana*

**DOI:** 10.1101/2024.03.04.582885

**Authors:** Qian Ma, Sijia Liu, Sara Raggi, Siamsa M. Doyle, Barbora Pařízková, Deepak Kumar Barange, Edward G. Wilkinson, Isidro Crespo Garcia, Joakim Bygdell, Gunnar Wingsle, Dirk Roeland Boer, Lucia C. Strader, Fredrik Almqvist, Ondřej Novák, Stéphanie Robert

**Author notes:** Q.M. and S.L. contributed equally to this work. Department of Plant Biotechnology and Bioinformatics, Ghent University, and Center for Plant Systems Biology, VIB, 9052, Ghent, Belgium. College of Horticulture, Shenyang Agricultural University, Shenyang, 110866, China (Sijia Liu). Umeå Plant Science Centre, Department of Forest Genetics and Plant Physiology, Swedish University of Agricultural Sciences, SE-901 83 Umeå, Sweden. **Author Contributions:** Q.M., S.L., and S.Ro. designed research; Q.M., S.L., S.Ra., S.M.D., B.P., D.K.B., E.G.W., I.C.G., J.B., G.W., and D.R.B. performed research and analyzed data; I.C.G., and D.R.B. contributed new reagents/analytical tools; L.C.S., F.A., O.N., and S.Ro. supervised research; S.L., S.M.D., and S.Ro. wrote the paper.

## Abstract

Apical hook development is an ideal model for studying differential growth in plants, and is controlled by complex hormonal crosstalk, with auxin and ethylene being the major players. Here, we identified a bioactive small molecule that decelerates apical hook opening in *Arabidopsis thaliana*. Our genetic studies suggest that this molecule enhances or maintains the auxin maximum found in the inner hook side and requires certain auxin and ethylene signaling components to modulate apical hook opening. Using biochemical approaches, we then revealed the WD40 repeat scaffold protein RECEPTOR FOR ACTIVATED C KINASE 1A (RACK1A) as a direct target of this compound. We present data in support of RACK1A playing a positive role in apical hook opening by negatively regulating the differential auxin response gradient across the hook via specific auxin and ethylene signaling mechanisms and thereby adjusting differential cell growth, an essential process for organ structure and function in plants. We have thus identified a role for RACK1A and auxin-ethylene crosstalk in negatively regulating differential cell growth to promote apical hook opening.

**Significance Statement:** Differential growth, or the growth of cells at different rates across tissues, is essential for providing shape and structure during plant development. The apical hook is a transient structure formed by differential cell growth across the hypocotyl tip in dark-grown seedlings, which protects the underlying tissues, and which opens during seedling development. We identified a small molecule that decelerates hook opening and discovered that it targets the protein RECEPTOR FOR ACTIVATED C KINASE 1A (RACK1A). We then showed that RACK1A promotes apical hook opening at the level of crosstalk between the plant hormones auxin and ethylene, by adjusting differential cell growth. Our work paves the way to a better understanding of how plants regulate and adapt their growth during development.

## Introduction

In the natural environment, plants germinate and start developing in the dark surrounded by soil, from which they absorb moisture and nutrients. Dicotyledonous plants have evolved the apical hook, a transient structure at the hypocotyl apex that preserves the integrity of the shoot apical meristem during the early stages of development (Zadnikova et al., 2016). The timing of the apical hook developmental process has been well characterized in *Arabidopsis thaliana* and three phases, named formation, maintenance, and opening phases, can be distinguished (Raz and Ecker, 1999). The formation phase begins when hypocotyl bending starts and proceeds until maximum curvature is reached. During this phase, hook curvature is generated by differential cell elongation on either side of the hypocotyl as it grows, with the cells in the outer side of the hook elongating more than those in the inner side. The hook remains at its maximum level of curvature at the growing hypocotyl apex during the maintenance phase. Finally, the hypocotyl apex straightens, and the hook opens during the opening phase. During this phase, the cell elongation rate in the inner side of the hook increases until cell length is similar on both sides of the hypocotyl apex (Silk and Erickson, 1978; Raz and Ecker, 1999).

Apical hook development is tightly regulated by multiple plant hormones, of which auxin and ethylene have been widely studied (Abbas et al., 2013; Mazzella et al., 2014; Wang and Guo, 2019). An asymmetrical auxin response has been observed across the inner and outer sides of the hook, with an auxin response maximum present in the inner side (Li et al., 2004). This auxin response gradient has been proposed to determine the differential cell elongation on the two sides of the hypocotyl apex that lead to hook formation and maintenance, with the high auxin response in the inner hook side repressing cell growth (Béziat and Kleine-Vehn, 2018; Wang and Guo, 2019). Similarly, loss of this auxin response maximum at the late maintenance phase releases cell growth repression in the inner hook side, leading to hook opening. Disruption of auxin synthesis, transport or signaling through genetic mutations or pharmacological approaches consequently leads to impaired apical hook development. For instance, seedlings harboring a *YUCCA*-dominant mutation and mutants defective in both *TRYPTOPHAN AMINOTRANSFERASE OF ARABIDOPSIS 1* (*TAA1*) and *TAA RELATED 2* (*TAR2*), which have increased or decreased auxin biosynthesis respectively, show abolished or reduced hook curvature (Zhao et al., 2001; Stepanova et al., 2008; Vandenbussche et al., 2010; Cao et al., 2019). Upon exogenous auxin or auxin efflux inhibitor 1-naphthylphthalamic acid (NPA) treatment, etiolated seedlings are hookless, supporting the necessity of establishing an auxin gradient for normal hook formation (Žádníková et al., 2010). In addition, mutations in the auxin perception gene families *TRANSPORT INHIBITOR RESPONSE 1* (*TIR1*)*/AUXIN BINDING F BOX PROTEINS* (*ABFs*) and *AUXIN/INDOLE-3-ACETIC ACID* (*AUX/IAAs*), such as in the gain-of-function mutants *auxin resistant 5* (*axr5)/iaa1* and *axr3/iaa17*, as well as in auxin signaling genes, such as in *axr1* and the loss-of-function mutant *nonphototropic hypocotyl 4* (*nph4*; also known as *auxin response factor 7* (*arf7*)), lead to severe hook defects (Leyser et al., 1993; Harper et al., 2000; Dharmasiri et al., 2005; Žádníková et al., 2010; Vain et al., 2019).

On the other hand, ethylene has been reported to enhance apical hook curvature. Etiolated *Arabidopsis* seedlings treated with exogenous ethylene, and *ethylene overproduction* (*eto*) and *constitutive ethylene-response 1* (*ctr1*) mutants with a constitutively activated ethylene signaling cascade, exhibit exaggerated apical hooks (Bleecker et al., 1988; Guzmán and Ecker, 1990; Vandenbussche et al., 2010; An et al., 2012). Kinematic analysis revealed that this feature is caused by a delay in the transition between formation and maintenance phases (Vandenbussche et al., 2010; Žádníková et al., 2010; Jonsson et al., 2017). In contrast, ethylene-insensitive mutants, such as *ethylene resistant 1* (*etr1*), *ethylene insensitive 2* (*ein2*), and double *ethylene insensitive 3* (*ein3*) *ein3-like 1* (*eil1*) show reduced hook curvature (Bleecker et al., 1988; Guzmán and Ecker, 1990; An et al., 2012). Extensive evidence supports the importance of crosstalk between ethylene and auxin in the regulation of apical hook development, with *HOOKLESS 1* (*HLS1*) being a key integrator between the two signaling pathways (Van de Poel et al., 2015; Béziat and Kleine-Vehn, 2018; Wang and Guo, 2019). Specifically, ethylene promotes apical hook curvature by stimulating the expression of *HLS1*, a direct downstream target of EIN3 and EIL1, which suppresses the expression of the auxin signaling repressor gene *ARF2*, thereby increasing the auxin response (Lehman et al., 1996; Li et al., 2004; An et al., 2012). Moreover, the reduced apical hook curvature observed in ethylene-insensitive mutants *etr1*, *ein2*, *ein3* and *ein4* can be fully restored when low concentration of auxin is applied, suggesting that ethylene is necessary to achieve the appropriate auxin level for normal hook formation (Vandenbussche et al., 2010). In addition, ethylene has also been shown to activate the expression of the auxin biosynthesis gene *TAR2*, auxin influx carrier gene *AUXIN-RESISTANT 1* (*AUX1*), and auxin efflux carrier genes *PIN-FORMED 3* (*PIN3*) and *PIN7* during apical hook development (Vandenbussche et al., 2010; Žádníková et al., 2010). However, many details of the roles of ethylene-auxin crosstalk in apical hook development still remain to be unraveled.

The ubiquitin-proteasome system has been reported to play a prominent regulatory role in plant hormone signaling (Santner and Estelle, 2010; Kelley and Estelle, 2012). The ubiquitin-like protein RELATED TO UBIQUITIN/NEURAL PRECURSOR CELL EXPRESSED DEVELOPMENTALLY DOWN-REGULATED PROTEIN 8 (RUB/NEDD8) uses similar enzymatic machineries as ubiquitin for NEDD8 conjugation (neddylation) (Schwechheimer, 2018). AUXIN RESISTANT 1 (AXR1) is a subunit of a heterodimeric NEDD8-activating enzyme (del Pozo et al., 1998), and mutation in the *AXR1* gene impairs sensitivity to multiple phytohormones, including auxin (Leyser et al., 1993; del Pozo et al., 2002; Tiryaki and Staswick, 2002; Vandenbussche et al., 2010).

Here we applied a chemical biology screen to identify novel bioactive small molecules that target plant development, such as we have done previously (Vain et al., 2019), resulting in isolation of the compound DAPIA (<u>D</u>elay of <u>Ap</u>ical Hook Opening in *<u>a</u>xr1-30*). By combining molecular genetics and biochemical approaches, we pinpoint a WD40 repeat scaffold protein, RECEPTOR FOR ACTIVATED C KINASE 1A (RACK1A), as the direct target of DAPIA and provide evidence that RACK1A is a component involved in the auxin/ethylene crosstalk regulating the mechanisms governing differential cell growth during apical hook opening.

## Results

### Identification of the small molecule DAPIA, which affects apical hook development in *Arabidopsis thaliana*

We previously developed a screening strategy for the identification of compounds targeting differential plant growth and development, based on difference in sensitivity between *Arabidopsis thaliana* Columbia-0 (Col-0) wild-type (WT) and the multiple phytohormone-insensitive mutant *axr1-30* (Vain et al., 2019). We used a similar strategy in the present work. From a screen of 4560 diverse chemicals, we obtained a small molecule that we named DAPIA (Fig. 1A), which was active in rescuing apical hook development in *axr1-30* in a dose-dependent manner, without obviously affecting the apical hook angle of WT, after 4 days of growth in darkness (Fig. 1B and C). Reduced hypocotyl length was also noted in the etiolated seedlings grown in the presence of the compound, for both WT and *axr1-30* (Fig. 1B).

**Fig. 1.**
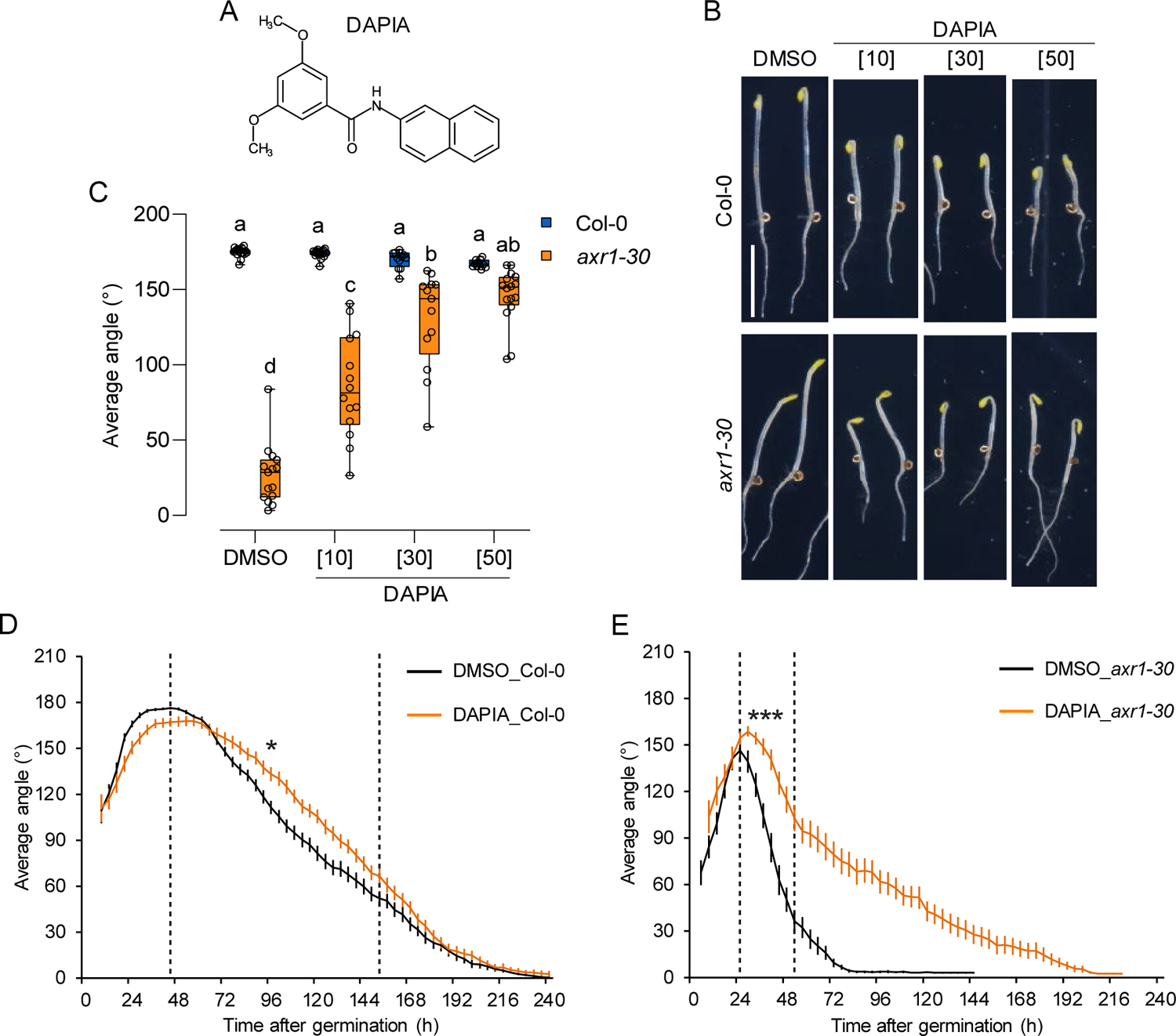
The compound DAPIA decelerates apical hook opening in *Arabidopsis* etiolated seedlings. (*A*) Chemical structure of DAPIA. (*B* and *C*) DAPIA dose-response analysis – representative images of apical hook phenotypes (*B*) and quantification of average hook angle (*C*) after 4 days of growth of Col-0 and *axr1-30* in darkness on medium supplemented with DMSO (mock) or DAPIA. Values in square brackets represent concentrations in µM. Scale bar represents 5mm. Data is shown as box plots and letters indicate significantly different means of n = 8-17 seedlings at *P* < 0.05 (one-way ANOVA, Tukey’s multiple comparisons test) (*C*). (*D* and *E*) Kinematics of apical hook angle in Col-0 (*D*) and *axr1-30* (*E*) as measured every 4 hours for 10 days of growth starting from germination (0 h) in darkness on medium supplemented with DMSO (mock) or 10 µM DAPIA. Error bars represent SEM; n = 23-61 seedlings. Dashed lines indicate the late maintenance-opening phase of the mock-treated control, for which asterisks indicate significantly different kinematic curves (CGGC method; \**P* < 0.05; \*\*\**P* < 0.001).

To further clarify the mode of action of DAPIA on apical hook development, we performed a kinematic analysis by continuously recording the apical hook angle of etiolated Col-0 and *axr1-30* seedlings, as previously described (Vandenbussche et al., 2010). We observed that DAPIA did have an effect in the WT, delaying the formation phase as well as reducing the maximum hook angle and decelerating hook opening (*SI Appendix*, Fig. S1*A*). As the mechanisms regulating the formation and opening phases of apical hook development are distinct (Béziat et al., 2017), we chose to focus on the late maintenance-opening phase, during which differential growth is repressed, and equal growth is restored, across the apical hook sides. We performed statistical comparison of the mock- and DAPIA-treated data at the time points representing this phase in the control (see Methods), revealing a significant effect of the compound on the late maintenance-opening phase in the WT (Fig. 1D). Regarding the *axr1-30* mutant, it displayed a smaller maximum hook angle than the WT, no maintenance phase and much more rapid hook opening than the WT in control conditions (*SI Appendix*, Fig. S1*A*). DAPIA increased the maximum hook angle in *axr1-30* as well as strongly decelerating hook opening (*SI Appendix*, Fig. S1*A*) and this effect of DAPIA in the opening phase was highly significant (Fig. 1E). Since germination takes approximately 24-36 hours in our hands, the DAPIA dose response data presented earlier at 4 days of growth (Fig. 1B and C) corresponds to about 60-72 hours after germination, at which time the WT is barely affected, while *axr1-30* is strongly affected, by DAPIA treatment (*SI Appendix*, Fig. S1*A*). Together, these results suggest that the compound DAPIA targets a pathway promoting apical hook opening in the WT and that *axr1-30* may be more sensitive to this effect. As AXR1 lies upstream of multiple hormone signaling pathways, we speculate that defects in other non-DAPIA-targeted pathways regulating hook opening in *axr1-30* may lead to stronger DAPIA-induced repression of hook opening than in the WT.

To gain insight into DAPIA activity, we performed a structure-activity relationship (SAR) study. The chemical structure of DAPIA suggests possible cleavage, which would release an amide (DAPIA-N) or a carboxylic acid (DAPIA-C) (*SI Appendix*, Fig. S1*B*). These two potential metabolites, as well as several DAPIA analogs that we synthesized, were assayed for their capacity to affect apical hook development in dark conditions (*SI Appendix*, Fig. S1*B* to *D*). 10 µM treatment with the potential metabolites DAPIA-N or DAPIA-C had no effect on apical hook angles of Col-0 or *axr1-30* after 4 days of dark growth (*SI Appendix*, Fig. S1*C*), suggesting the typical physiological effects shown by DAPIA are caused by the intact molecule rather than its metabolites. Similarly, none of the analogs tested exhibited an effect like that of DAPIA in rescuing *axr1-30* hook angle after 4 days, with most analogs having no effect at all, and only one analog, DAPIA-06, having a slight effect (*SI Appendix*, Fig. S1*D*). This suggests that DAPIA activity is linked to the whole molecule structure, as removals or modifications at various positions of the molecule generally abolish its bioactivity.

Due to the presence of a peptide bond (–CO–NH–) in the DAPIA structure, the compound was analyzed for possible metabolic conversion, with DAPIA-N and DAPIA-C being the most likely metabolites. The chemical stability of DAPIA was tested by analyzing the presence of DAPIA-N and DAPIA-C in the growth medium and *in planta*. In growth medium containing DAPIA, neither of the two potential metabolites was detectable with or without the growth of seedlings on the medium (*SI Appendix*, Fig. S2*A*). Without plants, the concentration of DAPIA in the medium remained unchanged after treatment for 4 days. However, the presence of growing seedlings resulted in around a 50% reduction in the concentration of DAPIA in the medium, suggesting that DAPIA is efficiently taken up by the plants (*SI Appendix*, Fig. S2*A*). Only DAPIA and negligible concentrations of DAPIA-N (0.004% and 0.005% of total DAPIA concentration in Col-0 and in *axr1-30,* respectively) were detectable in seedlings grown in the presence of DAPIA for 4 days (*SI Appendix*, Fig. S2*A* to *C*). Overall, these results imply that DAPIA is chemically stable both *in vitro* and *in planta*.

### DAPIA decelerates apical hook opening and requires the auxin signaling components ARF2, ARF7 and ARF19 for this effect

The asymmetric auxin response that is essential for apical hook development includes an auxin response maximum in the inner side of the hook (Vandenbussche et al., 2010). In contrast to the effects of DAPIA on apical hook development (Fig. 1D), exogenous auxin treatment abolishes hook formation in dark-grown seedlings (Žádníková et al., 2010), suggesting that DAPIA is physiologically distinct from auxin. However, we found that the application of DAPIA resulted in an enlarged region of auxin response maximum in the inner side of the hook of the WT after 4 days of growth in darkness, as observed via *DR5::GUS* auxin response reporter expression (Fig. 2A). Thus, DAPIA may exert its effect on decelerating hook opening via locally enhancing the auxin response maximum in the inner side of the hook. Alternatively, since 4 days of growth corresponds to about 60-72 hours after germination as described earlier, DAPIA may rather maintain the auxin response maximum in the inner side, which should be starting to reduce in the untreated control at this late maintenance-opening phase time point (see Fig. 1A).

**Fig. 2.**
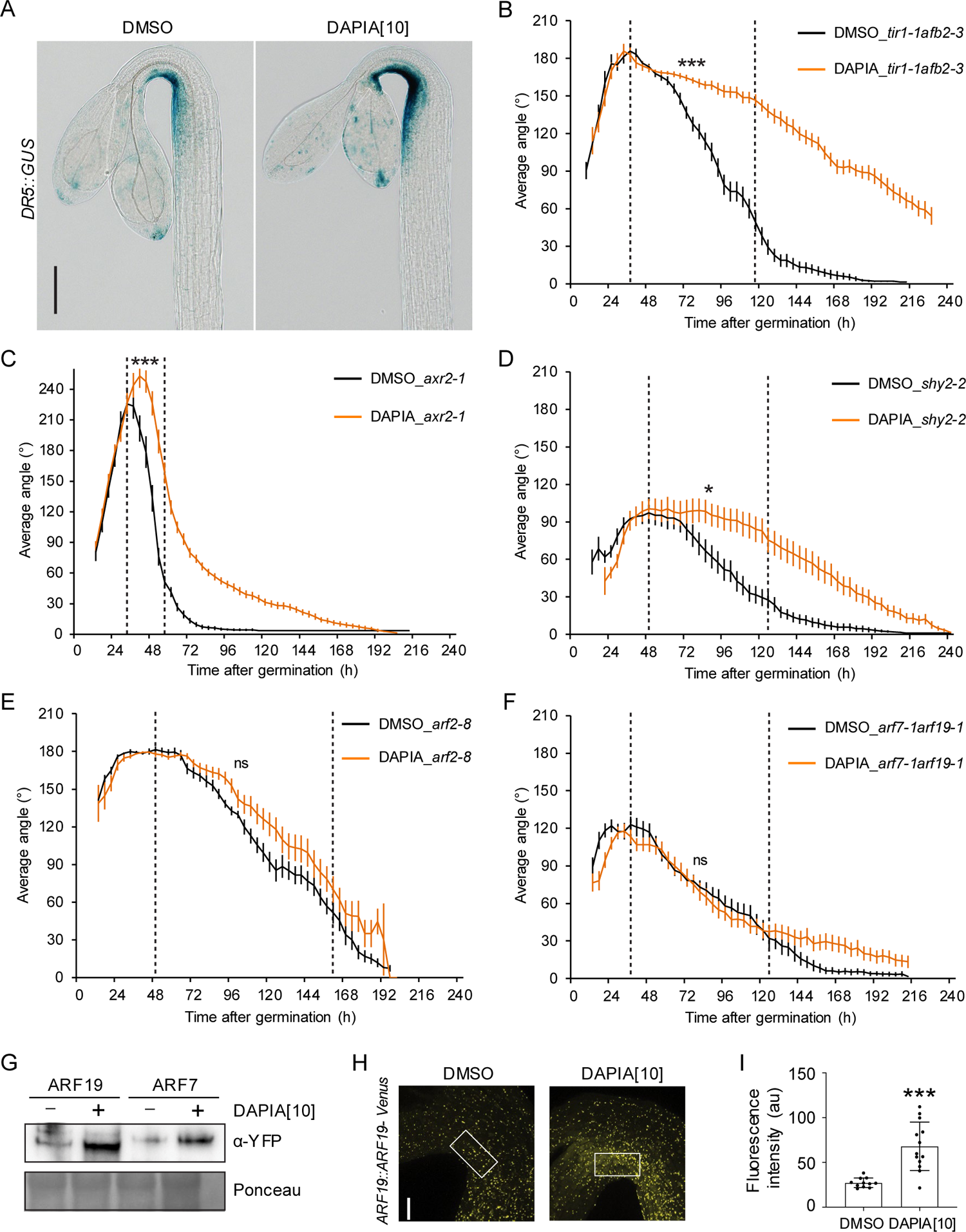
DAPIA affects the auxin response in the hook and requires auxin signaling components ARF2, ARF7 and ARF19 to decelerate hook opening. (*A*) Representative images of apical hooks of GUS-stained *DR5::GUS* seedlings after 4 days of growth in darkness on medium supplemented with DMSO (mock) or DAPIA. Values in square brackets represent concentrations in µM. Scale bar represents 150 µm. (*B* to *F*) Kinematics of apical hook angle in *tir1-1afb2-3* (*B*), *axr2-1* (*C*), *shy2-2* (*D*), *arf2-8* (*E*) and *arf7-1arf19-1* (*F*) as measured every 4 hours for 10 days of growth starting from germination (0 h) in darkness on medium supplemented with DMSO (mock) or 10 µM DAPIA. Error bars represent SEM; n = 15-38 seedlings. Dashed lines indicate the late maintenance-opening phase of the mock-treated control, for which asterisks indicate significantly different kinematic curves (CGGC method; ns – not significantly different; \**P* < 0.05; \*\*\**P* < 0.001). (*G*) Immunoblot analysis of tissue samples from *ARF19:ARF19-Venus* and *ARF7:ARF7-Venus* seedlings, grown for 4 days in darkness on medium supplemented with DMSO (mock) or 10 µM DAPIA, probed with anti-YFP antibody. Ponceau stain was used as loading control. (*H* and *I*) Representative confocal images (*H*) and quantification of Venus signal intensity on inner side (*I*) of apical hooks of *ARF19::ARF19-Venus* seedlings after 4 days of growth in darkness on medium supplemented with DMSO (mock) or 10 µM DAPIA. Scale bar represents 50 µm. White boxes in (*H*) represent areas quantified in (*I*). Images of maximum intensity projections of z-stacks of Venus channel are shown. Graph represents one replication of three similar experiments, error bars represent SD, and asterisks indicate significantly different means (Student’s T-test; \*\*\**P* < 0.001). Values in square brackets represent concentrations in µM.

To investigate possible involvement of auxin signaling in DAPIA activity, we performed hook angle kinematic analysis in additional auxin perception-defective and various auxin signaling-defective mutants in the absence or presence of DAPIA. Compared to the Col-0 WT, the auxin receptor double mutant *tir1-1afb2-3* exhibited a slightly exaggerated maximum hook angle, followed by earlier, more rapid hook opening (*SI Appendix*, Fig. S3*A*). The application of DAPIA, however, strongly decelerated hook opening in *tir1-1afb2-3*, resulting in a highly significant effect of the compound on the late maintenance-opening phase (Fig. 2B). This over-sensitivity of *tir1-1afb2-3* to the effect of DAPIA on hook opening, like for *axr1-30*, may possibly be explained by the upstream role of the TIR1/AFB receptors in multiple auxin signaling pathways.

The AUX/IAA gain-of-function mutant *axr2-1/iaa7* exhibited an exaggerated hook, no maintenance phase and much faster hook opening than the Col-0 WT in control conditions (*SI Appendix*, Fig. S3*B*). The main effect of DAPIA in *axr2-1* was to enhance the already exaggerated hook, leading to a significant effect on the opening phase (Fig. 2C). Much like Col-0, hook formation was slightly delayed, maximum hook angle was decreased and hook opening was somewhat decelerated by DAPIA treatment in the Landsberg *erecta* (Ler) WT, leading to a significant effect on the late maintenance-opening phase (*SI Appendix*, Fig. S3*C*). The AUX/IAA gain-of-function mutant *shy2-2/iaa3* exhibited a smaller maximum hook angle than the Ler WT as well as slower hook opening in control conditions (*SI Appendix*, Fig. S3*D*). DAPIA treatment had a significant effect on *shy2-2*, extending the maintenance phase as well as further decelerating hook opening (Fig. 2D). We next tested the activity of DAPIA on *arf2-8*, a loss of function mutant in the transcriptional modulator ARF2 shown to be involved in apical hook development through the ethylene signaling pathway (Li et al., 2004), which displayed similar hook angle kinetics to the Col-0 WT in control conditions (*SI Appendix*, Fig. S3*E*). Although DAPIA treatment appeared to slightly decelerate hook opening in *arf2-8*, the effect on the late maintenance-opening phase was not statistically significant, indicating resistance of this mutant to the effect of DAPIA on hook opening (Fig. 2E). Finally, *arf7-1arf19-1*, a loss-of-function double mutant of the auxin transcriptional activators ARF7 and ARF19, which showed a strong impairment in hook formation and somewhat slower hook opening compared to the Col-0 WT in control conditions (*SI Appendix*, Fig. S3*F*), was also resistant to the effects of DAPIA on the late maintenance-opening phase (Fig. 2F). Taken together, these results show that DAPIA requires the functional auxin signaling components ARF2, ARF7 and ARF19 to exert its effect on decelerating apical hook opening.

Since *arf7-1arf19-1* displays somewhat slower hook opening than the WT and ARF7 and ARF19 are required for DAPIA to exert its effect on decelerating apical hook opening, these transcription factors seem likely to play a role in the opening process. We were curious as to what effect DAPIA might have on the abundance of ARF7 and ARF19 at the protein level. We therefore analyzed tissue samples from *ARF7::ARF7-Venus* and *ARF19::ARF19-Venus* etiolated seedlings grown in the presence of DAPIA for abundance of the Venus protein by Western blotting, revealing a strong increase in protein abundance compared to the mock control at the whole seedling level (Fig. 2G). To investigate this effect specifically in the apical hook, we performed confocal microscopy imaging of etiolated *ARF19::ARF19-Venus* seedling hooks, revealing a clear DAPIA-induced enhancement of the fluorescence signal (Fig. 2H). In particular, DAPIA induced an enhanced cytosol-to-nuclear fluorescence signal in the inner hook side. Quantification of the overall fluorescence signal at the inner hook side confirmed that DAPIA enhances ARF19-Venus protein abundance in this region (Fig. 2I), which may be a means by which DAPIA enhances or maintains an auxin response maximum in the inner side of the apical hook and consequently disrupts hook opening.

### The deceleration of apical hook opening by DAPIA requires ethylene signaling components

Ethylene is known to play a positive role in apical hook formation (Lehman et al., 1996), for which its crosstalk with auxin is thought to be important (Li et al., 2004; Stepanova et al., 2007, 2008). We therefore questioned whether DAPIA might modulate apical hook opening through the ethylene biosynthesis or signaling pathways. Exogenous treatment with ethylene or the ethylene biosynthesis precursor 1-aminocyclopropane-1-carboxylic acid (ACC) results in exaggerated apical hook formation in etiolated seedlings (Guzmán and Ecker, 1990; An et al., 2012; Vandenbussche et al., 2010), an effect that is not mimicked by DAPIA (Fig. 1D), suggesting that DAPIA is physiologically distinct from ethylene. We therefore turned our focus to the ethylene signaling pathway. The ethylene-insensitive mutants *ein2-t*, *ein3-1eil1-1*, *etr1-1* and *hls1-1* were selected for hook angle kinematic analysis in the presence of DAPIA. In control conditions, all these mutants exhibited various degrees of defects in hook development compared to the Col-0 WT, including a smaller maximum angle of hook curvature, a shorter or absent maintenance phase and more rapid hook opening (*SI Appendix*, Fig. S4*A* to *D*). Interestingly, the ethylene perception mutants *ein2-t* and *ein3-1eil1-1* and signaling mutant *hls1-1* all showed resistance to the effects of DAPIA on the late maintenance-opening phase, while the ethylene signaling mutant *etr1-1* was significantly affected by DAPIA at this phase (Fig. 3A to D). In particular, *ein2-t* and *hls1-1* were totally unaffected by DAPIA treatment (Fig. 3A and D). Although there appeared to be a slight effect of DAPIA on both *ein3-1eil1-1* and *etr1-1*, only *etr1-1* showed a significant difference to the WT at the late maintenance-opening phase (Fig. 3B and C). These results suggest that DAPIA requires certain ethylene signaling components for modulating the apical hook opening process.

**Fig 3.**
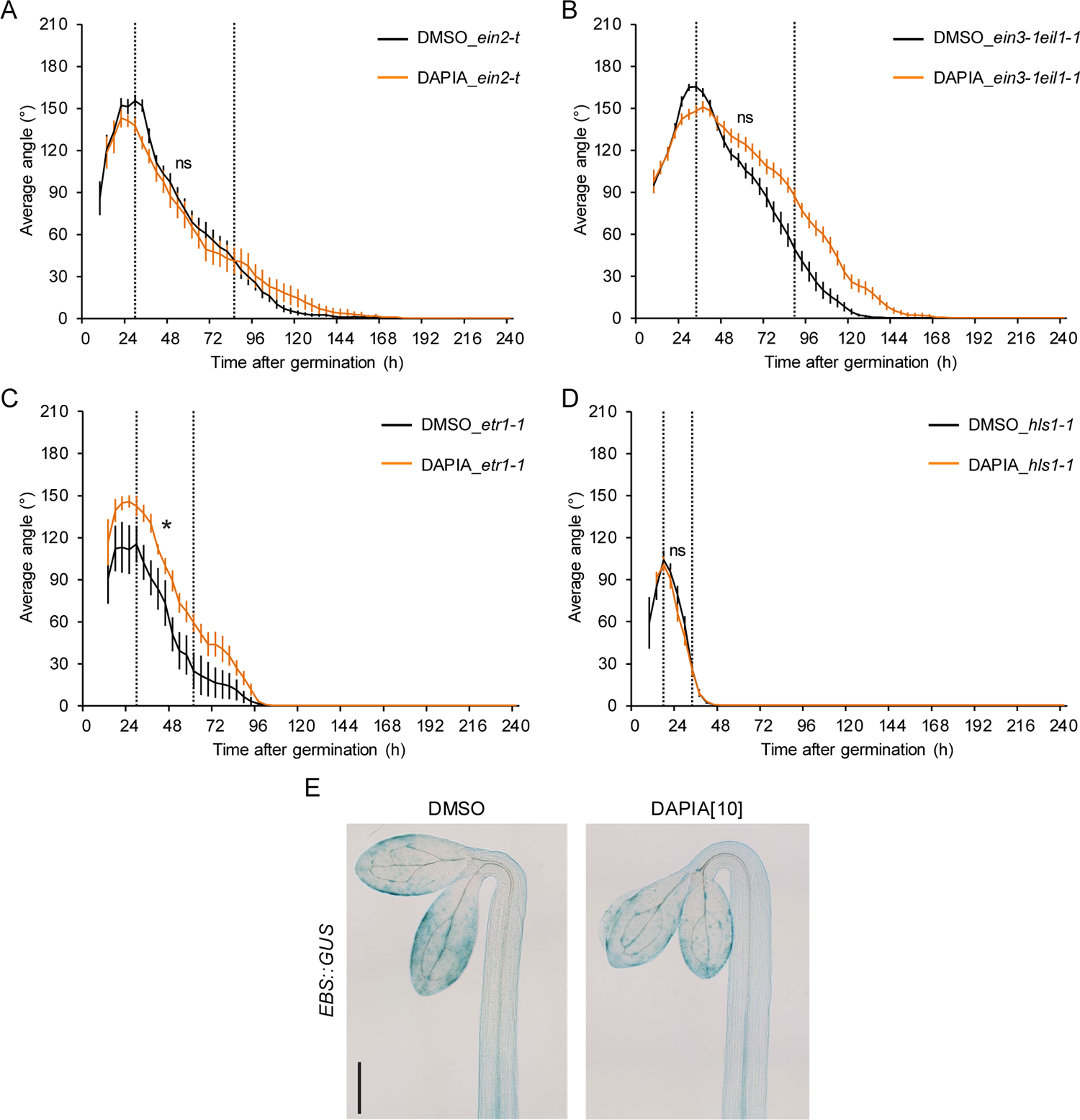
DAPIA requires ethylene signaling components to decelerate hook opening. (*A* to *D*) Kinematics of apical hook angle in *ein2-t* (*A*), *ein3-1eil1-1* (*B*), *etr1-1* (*C*) and *hls1-1* (*D*) as measured every 4 hours for 10 days of growth starting from germination (0 h) in darkness on medium supplemented with DMSO (mock) or 10 DAPIA. Error bars represent SEM; n = 7-30 seedlings. Dashed lines indicate the late maintenance-opening phase of the mock-treated control, for which asterisks indicate significantly different kinematic curves (CGGC method; ns – not significantly different; \**P* < 0.05). (*E*) Representative images of apical hooks of GUS-stained *EBS::GUS* seedlings after 4 days of growth in darkness on medium supplemented with DMSO (mock) or DAPIA. Values in square brackets represent concentrations in µM. Scale bar represents 150 µm.

Ethylene signaling has been proposed to modulate auxin transport, leading to a dynamic asymmetric auxin response that is essential for apical hook development (Vandenbussche et al., 2010; Žádníková et al., 2010). Our data suggest that DAPIA acts via ethylene signaling components to decelerate hook opening, which led us to examine the effects of DAPIA on ethylene responses in etiolated seedlings at the transition between the hook maintenance and opening phases. However, DAPIA treatment did not alter the very faint ethylene response signal observed in the hook of the *EBS::GUS* ethylene response reporter (Fig. 3E).

We also analyzed hook angle kinematics in the gibberellin-insensitive mutant *gai-1*, in the *wag2-1* mutant defective in a kinase shown to repress hook opening downstream of gibberellin (Willige et al., 2012), and in the cytokinin signaling double mutant *ahk2-2ahk3-3*. The *gai-1* mutant presented a smaller maximum hook angle than the Ler WT as well as a much shorter maintenance phase (*SI Appendix*, Fig. S5*A*) and was significantly sensitive to the effect of DAPIA on the late maintenance-opening phase (*SI Appendix*, Fig. S5*B*). In contrast, *wag2-1* showed more rapid hook formation and opening than the Col-0 WT (*SI Appendix*, Fig. S5*C*) but was also significantly sensitive to the DAPIA effect (*SI Appendix*, Fig. S5*D*). Finally, *ahk2-2ahk3-3* presented an extended maintenance phase and a more rapid rate of hook opening than the WT (*SI Appendix*, Fig. S5*E*) and was also sensitive to the effect of DAPIA on the late maintenance-opening phase (*SI Appendix*, Fig. S5*F*). These results suggest that DAPIA may not affect apical hook opening via gibberellin or cytokinin signaling pathways.

### RACK1A is a direct target of DAPIA

To identify the protein target of DAPIA, we performed a *de novo* drug affinity responsive target stability (DARTS) assay (Pai et al., 2015), a technology based on the evidence that some proteins, when bound to chemical ligands, are protected from degradation by proteases. We reasoned that the stronger effect of DAPIA on decelerating hook opening in *axr1-30* compared to the WT (Fig. 1D and E) might enable easier identification of the protein target of DAPIA in the mutant. We thus incubated total protein extracts from *axr1-30* etiolated seedlings with DAPIA and treated with the proteolytic enzyme pronase before subsequent analysis by LC-MS/MS, leading to the identification of several protein candidates as being protected from digestion and therefore potential targets of DAPIA (*SI Appendix*, Dataset S1). We chose to focus on one particular candidate, RACK1A, as it is known to be involved in multiple hormone signaling pathways (Islas-Flores et al., 2015), but has not previously been implicated in the regulation apical hook development. This protein is the *Arabidopsis* homologue of the tobacco WD40 repeat ArcA, described as a scaffold protein involved in glucose, gibberellin and abscisic acid signaling pathways (Vahlkamp and Palme, 1997; Islas-Flores et al., 2015). To validate the binding ability of DAPIA with RACK1A, we tested the effects of DAPIA on pronase proteolytic degradation of RACK1A protein by immunoblotting in the *axr1-30* protein extracts (Fig. 4A). DAPIA started to show protection of RACK1A against proteolysis when the pronase:protein ratios were in the range of 1:1000 to 1:100, with the highest pronase level displaying the most obvious difference between DAPIA-treated and non-treated samples (Fig. 4A). These results provide support for an interaction between DAPIA and RACK1A *in vitro*.

**Fig. 4.**
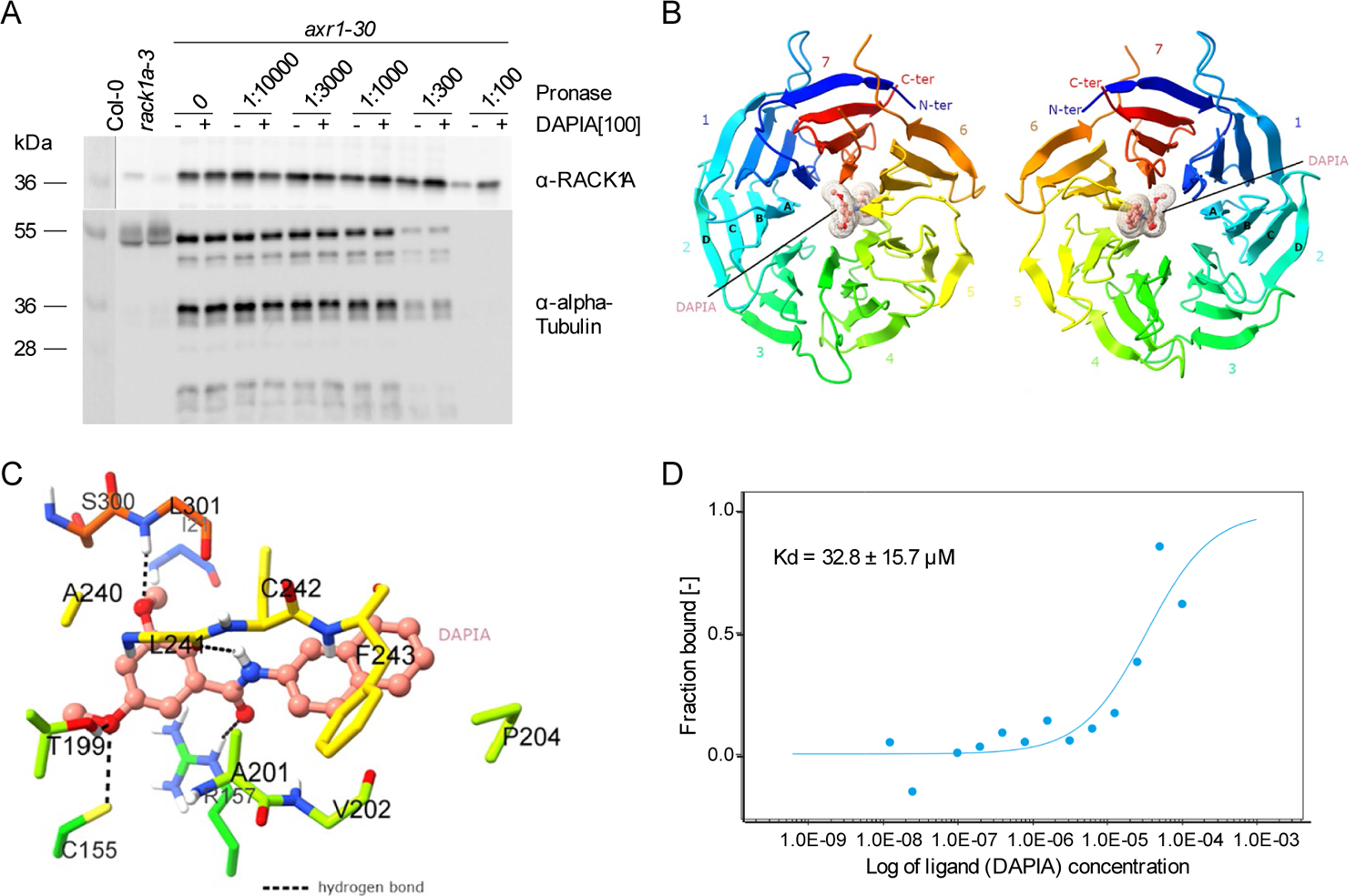
RACK1A is the direct target of DAPIA. (*A* to *D*) Direct binding between DAPIA and RACK1A validated by proteolytic analysis (*A*), molecular docking (*B* and *C*) and micro-scale thermophoresis (MST) analysis (*D*). (*A*) Total protein extracts from 4-day-old etiolated seedlings of *axr1-30*, as well as Col-0 and *rack1a-3* as controls, were incubated with DMSO (mock) or DAPIA and subjected to various dilutions of pronase (expressed as total enzyme:total protein ratios). RACK1A protein levels were detected through protein gel blot analysis using anti-RACK1A antibody and anti-alpha-Tubulin antibody as a reference protein control. Values in square brackets represent concentrations in µM. (*B*) Top (left panel) and bottom (right panel) views of RACK1A 3D structure with the DAPIA binding site predicted by the docking simulation. RACK1A, in ribbon diagram, adopts a β-propeller conformation. The seven blades of the β-propeller are numbered 1-7 from the N terminus (N-ter, blue) to the C terminus (C-ter, red) in rainbow mode. The four antiparallel β-strands in each blade are labelled as A-D, as exemplified in blade 2. The small molecule DAPIA is bound in a cavity in the central channel and is shown in stick-and-ball and surface mesh modes in salmon color. (*C*) A close-up view of DAPIA in the predicted binding pocket of RACK1A. The thirteen RACK1A residues involved in the interaction with DAPIA are shown in sticks, with the same color codes as used in b. Hydrogen bonds are shown as black dashed lines. (*D*) For MST analysis, Kd = mean ± SD of four independent replicates.

Next, we performed molecular docking analysis to explore the likely interaction between DAPIA and RACK1A and to predict the binding site. We docked DAPIA *in silico* to the available structure of *Arabidopsis* RACK1A (Protein Data Bank (PDB) code 3DM0) (Ullah et al., 2008). The overall protein structure adopts a seven-bladed β-propeller fold circularly arranged around a central channel with a diameter of ∼9Å, which is canonical for the WD40 family proteins (Ullah et al., 2008). Each propeller blade is composed of four antiparallel β-strands, named A to D (Fig. 4B). Dockings with DAPIA predicted only one specific binding site within the central channel of the β-propeller fold with a reasonable estimated free energy of binding of −9.28 kcal mol^-1^ (Fig. 4B). According to the interaction mode, DAPIA formed hydrogen bond interactions with Cys155, Arg157, Thr199, Leu241 and Leu301, and established extensive beneficial contacts with Ile21, Cys155, Arg157, Thr199, Ala201, Val202, Pro204, Ala240, Leu241, Cys242, Phe243, Ser300 and Leu301 (Fig. 4B and C). This simulation suggests that DAPIA may tightly bind to RACK1A via the binding pocket located in the protein’s central channel. The thirteen amino acid residues constituting the binding pocket are also conserved among *Arabidopsis*, human and *Drosophila*, with the exception of Ala201, which is replaced by Thr in the latter two species (*SI Appendix*, Fig. S6) (Ullah et al., 2008). Further analysis of the spatial distribution of these 13 amino acids revealed that they are all located at the inner β-strand A, involving blades 1 and 4 to 7, except for Pro204 that is positioned in the loop connecting strand A and B of blade 5. Moreover, eight of them are from blades 5 and 6 (four each). Previous studies revealed two conserved surface regions within the RACK1 proteins; conserved region 1 is located on the top face of the propeller and region 2 on the bottom. They impart a scaffolding function to the protein by mediating protein-protein interactions (Ullah et al., 2008). Conserved region 2 is mainly comprised of amino acids from blade 5 (Pro204, Asp205 and Tyr230) and blade 6 (Asn246, Tyr248 and Trp249) (Ullah et al., 2008). This result implies that β-propeller blades 5 and 6 may play an important role in facilitating interactions with the ligand and with other partner proteins. To summarize, our docking simulations suggest that DAPIA may target RACK1A through extensive interactions with a binding site buried in the central channel, which consequently may interfere with protein-protein interactions mediated by the conserved region 2 of RACK1A.

To confirm the docking result qualitatively and quantitatively, we characterized the specificity and quantified the kinetics and affinity of binding between DAPIA and RACK1A using microscale thermophoresis (MST). The inactive analogs DAPIA-02 and DAPIA-07 (see *SI Appendix*, Fig. S1*B* and *D*) were used as negative controls. In the absence of Tween 20 or PIC, the signal to noise (S/N) value of DAPIA-treated samples reached 46.9, implying good binding between DAPIA and RACK1A (*SI Appendix*, Fig. S7*A*), while the presence of Tween 20 or PIC abolished the binding (S/N value 1.4) (*SI Appendix*, Fig. S7*B*). The dose-response curves further revealed a dissociation constant (*K*_d_) of 32.8 ± 15.7 μM (SD), suggesting again the optimal binding affinity of DAPIA to RACK1A (Fig. 4D). Notably, the two inactive analogs, DAPIA-02 and DAPIA-07, did not show any binding affinity with RACK1A (*SI Appendix*, Fig. S7*C*). These results confirm that RACK1A is a direct target of DAPIA.

### RACK1A functions in apical hook opening at the level of crosstalk between auxin response and ethylene signaling

Our next step was to perform hook angle kinetic analysis in the loss-of-function mutant *rack1a-3*. In control conditions, *rack1a-3* displayed a similar maximum angle of hook curvature to the WT, but a longer maintenance phase and a much slower rate of early hook opening (*SI Appendix*, Fig. S8*A*). The difference between *rack1a-3* and Col-0 was highly significant at the late maintenance-opening phase (Fig. 5A). In fact, the effect of the *rack1a-3* mutation on the late maintenance-opening phase was stronger than that of DAPIA treatment in the WT (*SI Appendix*, Fig. S8*B*). These results support the idea that DAPIA decelerates hook opening by targeting RACK1A, suggesting that RACK1A is involved in regulating hook opening. Unlike the effect of DAPIA treatment in the WT control (*SI Appendix*, Fig. S8*B* and *C*), DAPIA treatment of *rack1a-3* resulted in a much smaller maximum angle of hook curvature and a shorter maintenance phase than in mock conditions (*SI Appendix*, Fig. S8*B*), leading to a highly significant effect on the late maintenance-opening phase (*SI Appendix*, Fig. S8*D*), although the rate of hook opening did not appear to be affected. These effects suggest that DAPIA may target alternative proteins involved in hook formation and maintenance, rather than opening, in the absence of RACK1A.

**Fig. 5.**
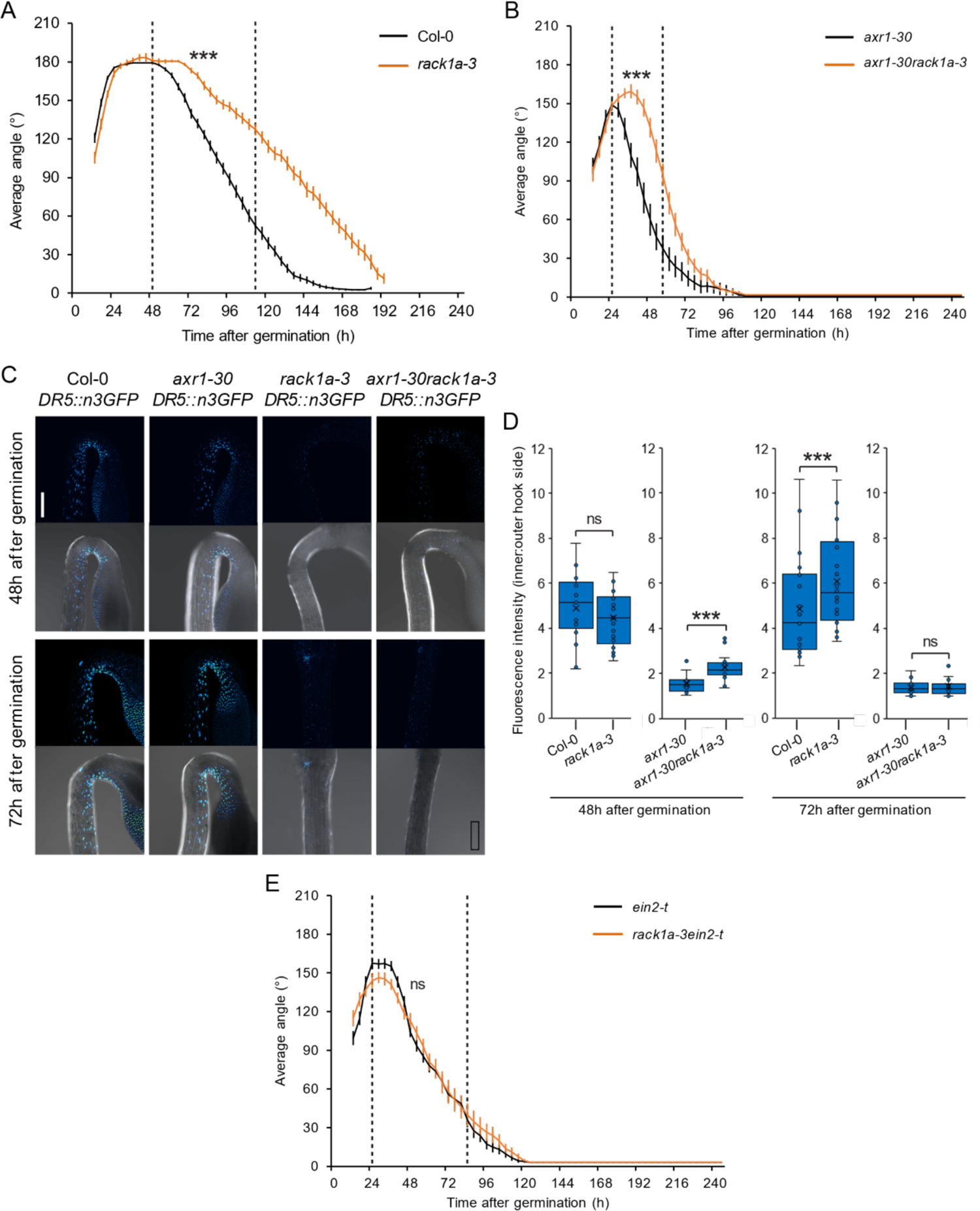
RACK1A regulates apical hook opening at the level of crosstalk between auxin response and ethylene signaling. (*A* and *B*) Kinematics of apical hook angle in Col-0 and *rack1a-3* (*A*) and in *axr1-30* and *axr1-30rack1a-3* (*B*) as measured every 4 hours for 10 days of growth starting from germination (0 h) in darkness. Error bars represent standard error of the mean; n = 22-26 seedlings. Dashed lines indicate the late maintenance-opening phase of Col-0 (*A*) or axr1-30 (*B*), for which asterisks indicate significantly different kinematic curves (CGGC method; \*\*\**P* < 0.001). (*C* and *D*) Representative confocal images (*C*) and quantification of auxin response gradient (inner:outer side average nuclear GFP signal intensity) (*D*) of apical hooks of *DR5::n3GFP* seedlings in Col-0, *axr1-30*, *rack1a-3* and *axr1-30rack1-a3* backgrounds 48 and 72 hours after germination and growth in darkness. Scale bar represents 200 µm. Low-to-high GFP signal intensity is represented as a blue-to-yellow color gradient, according to the inset. For GFP channels (top panels), maximum intensity projections of z-stacks with the same thickness and number of slices across samples are shown, and they are also shown merged with the brightfield images (bottom panels) (*C*). Data is shown as box plots and asterisks indicate significantly different means of n = 17-29 seedlings (Wilcoxon rank sum test; ns – not significantly different; \**P* < 0.05; \*\*\**P* < 0.001) (*D*). (*E*) Kinematics of apical hook angle in *ein2-t* and *rack1a-3ein2-t*, as for (*A*) and (*B*) with dashed lines indicating the late maintenance-opening phase of *ein2-t* (curves compared by CGGC method; ns – not significantly different).

In *Arabidopsis*, there are two other members of the *RACK1* gene family, *RACK1B* and *RACK1C*, which display high sequence similarity with *RACK1A* (Chen et al., 2006). We reasoned that RACK1B and RACK1C may be functionally redundant with RACK1A in the regulation of hook opening and therefore next analyzed hook angle kinetics in the estradiol-inducible triple RACK1A, B and C-targeted artificial microRNA mutant *amiR-rack1-es1*. Under estradiol treatment, *amiR-rack1-es1* displayed a delay in hook formation and opening compared to the WT (*SI Appendix*, Fig. S8*A*), which resulted in a highly significant difference to the WT at the late maintenance-opening phase (*SI Appendix*, Fig. S8*E*). However, the triple mutant was not more affected in this phase than *rack1a-3* (*SI Appendix*, Fig. S8*A* and *F*). We next investigated the effects of DAPIA on hook angle kinetics in the estradiol-induced triple mutant. The main effect was to decrease the maximum angle of hook curvature (*SI Appendix*, Fig. S8*G*), which led to a significant effect on the late maintenance-opening phase, but no change in the rate of hook opening (*SI Appendix*, Fig. S8*H*). These results suggest that RACK1B and RACK1C may not be involved in apical hook opening. Taken together, our results support the proposal that DAPIA decelerates apical hook opening through targeting of the RACK1A protein, implying a positive role of RACK1A in hook opening. As *rack1a-3* and triple *amiR-rack1-es1* showed some sensitivity to the effects of DAPIA treatment on the phases of hook formation and maintenance, there may also be other molecular pathways targeted by DAPIA as mentioned earlier, and this is supported by the identification of several other potential protein targets of this compound in our DARTS assay (*SI Appendix*, Dataset S1).

Since DAPIA partially rescues the early-opening apical hook phenotype of *axr1-30* (Fig. 1E), we hypothesized that mutations in RACK1A should achieve a similar result. To test this hypothesis, we crossed *axr1-30* and *rack1a-3* and performed hook angle kinetic analysis in the double mutant, revealing a distribution that was intermediate between that of the single mutants at the maintenance and opening phases (*SI Appendix*, Fig. S9*A*). Furthermore, statistical analysis confirmed that the *axr1-30* mutant was significantly rescued towards a WT-like hook angle kinetics phenotype at the late maintenance-opening phase by addition of the *rack1a-3* mutation (Fig. 5B).

Our results suggest that DAPIA can induce or maintain the auxin response in the apical hook at the end of the maintenance phase, resulting in deceleration of apical hook opening, and we hypothesized that mutations in RACK1A should achieve the same result. Since the early-opening apical hook phenotype of the *axr1-30* mutant is partially rescued by addition of the *rack1a-3* mutation (Fig. 5B), we also wondered what effect mutations in RACK1A would have on the auxin response in the hook of this mutant. To investigate this, we introduced the auxin response marker line *DR5::n3GFP* into the *rack1a-3* and *axr1-30rack1a-3* backgrounds. We examined the auxin response gradient across the apical hook by observing the *DR5::n3GFP* signal at both 48 and 72 hours after germination (time points at which hook opening is significantly affected in *axr1-30rack1a-3* compared to *axr1-30*, and in *rack1a-3* compared to the WT, respectively). At the 48 hours time point, there was no difference in the inner:outer side auxin response gradient across the apical hook between the WT and *rack1a-3* (Fig. 5C and D). The *axr1-30* mutant showed a much lower auxin response gradient across the hook than the WT, which agrees with its early hook opening phenotype. This low gradient was partially rescued at this time point by the introduction of *rack1a-3*, being slightly but significantly increased in *axr1-30rack1a-3* compared to *axr1-30* (Fig. 5C and D). This result agrees with the partial rescue of the *axr1-30* early hook opening phenotype at this time point by combination with *rack1a-3* that was shown earlier (Fig. 5B). At the 72 hours time point, *rack1a-3* displayed a significantly increased auxin response gradient compared to the WT, while the auxin response gradients in *axr1-30* and *axr1-30rack1a-3* were no longer different (Fig. 5C and D). These results support the idea that RACK1A positively regulates apical hook opening by suppressing the auxin response gradient in the hook.

According to our results, DAPIA can induce/maintain auxin but not ethylene responses in the apical hook at the late maintenance phase. However, the ethylene perception mutants *ein2-t* and *ein3-1eil1-1* and ethylene signaling defective mutant *hls1-1* showed resistance to the effect of DAPIA on the hook late maintenance-opening phase, leading us to hypothesize that DAPIA, and therefore RACK1A, may act at the crosstalk of auxin response and ethylene perception and/or signaling. We therefore next combined the *ein2-t* and *rack1a-3* mutations and performed hook angle kinetic analysis in the double mutant. As shown earlier, compared to the WT, *ein2-t* displayed accelerated hook opening (*SI Appendix*, Fig. S4*A*). Introduction of the *rack1a-3* mutation to the *ein2-t* mutant background resulted in loss of the *rack1a-3* decelerated hook opening phenotype (*SI Appendix*, Fig. S9*B*) and had no effect on the *ein2-t* hook angle kinetics curve at the late maintenance-opening phase (Fig. 5E), reminiscent of the complete resistance of *ein2-t* hook development to DAPIA treatment that we found previously (Fig. 3A). These results confirm that the effects of the compound DAPIA on the late maintenance-opening phase of apical hook development are a result of the targeting of RACK1A as well as supporting the hypothesis that RACK1A regulates apical hook opening via a mechanism requiring ethylene signaling.

Collectively, our results suggest that RACK1A plays a positive role in apical hook opening by negatively regulating the local auxin response maximum in the inner side of the hook via an ARF7/ARF19-regulated signaling pathway, at the level of crosstalk with ethylene signaling. In this way, RACK1A acts at the transition between the hook maintenance and opening phases to suppress differential growth and re-establish equal growth across both sides of the hook, inducing its opening.

## Discussion

Plant hormones and their crosstalk mediate the morphological adaptability and plasticity of plants that are essential for their survival and development. Although some crucial modulators involved in hormone signaling have been identified through genetic approaches (Ruegger et al., 1998; Guzmán and Ecker, 1990; Alonso et al., 2003; Li et al., 2004), many components remain unknown due to gene redundancy or loss-of-function lethality and complex interconnected crosstalk. The isolation of small, bioactive molecules has proven to be an effective approach towards the unraveling of complex hormone signaling pathways while overcoming many of these issues (De Rybel et al., 2009; Hayashi et al., 2008; Hayashi et al., 2012; Monte et al., 2014; Park et al., 2009; Rigal et al., 2021; Tsuchiya et al., 2010; Uchida et al., 2018; Vaidya et al., 2021; Vain et al., 2019). Despite many such effective approaches, few of them have reported the successful identification of the direct targets of the compounds. In this work, we isolated the small molecule DAPIA and unveiled its molecular target RACK1A as a modulator of apical hook opening at the level of crosstalk between auxin and ethylene signaling mechanisms. We employed DARTS in combination with proteomics analysis to pinpoint RACK1A, a WD repeat protein, as a direct target of DAPIA. Convincing evidence was further provided by an *in vitro* MST assay, showing that RACK1A displays an optimal binding affinity with DAPIA but not with its inactive analogs. Moreover, our *in vivo* data revealed that apical hook opening in the *rack1a-3* mutant is slower than in the WT, most likely as a consequence of enhanced or maintained inner:outer side auxin response gradient across the hook, which resembles the effects of DAPIA treatment in the WT, supporting RACK1A as a target of DAPIA. Among the three genes encoding RACK1 proteins in *Arabidopsis*, *RACK1A* shows the highest expression in various tissues and organs (Guo and Chen, 2008), implying its predominant role in the regulation of various developmental processes. Although RACK1A may redundantly regulate apical hook opening together with RACK1B and C, disruption of the single *RACK1A* gene is sufficient to decelerate hook opening, without any additive effect observed in a triple *RACK1* mutant, indicating *RACK1A* as the main regulator of this process.

Much of the work that has unraveled essential roles for auxin-ethylene crosstalk in apical hook development has been focused on the formation phase, during which the establishment of a precise auxin gradient across the hook is essential. Although hook opening requires subsequent reduction of this gradient, the signaling pathways regulating the appearance and disappearance of the auxin maximum appear to be distinct. For example, experimental evidence suggests that intercellular auxin transport mechanisms are involved in the establishment of the auxin maximum required for hook formation, while the depletion of the auxin maximum required for hook opening mainly involves intracellular alterations of auxin (Béziat et al., 2017). We therefore took advantage of the effect of DAPIA in decelerating apical hook opening to investigate the molecular events involving RACK1A that govern this poorly studied phase of hook development. Our reverse genetics approach, in which we tested auxin and ethylene perception and signaling mutants for their sensitivity to the effect of DAPIA on the late maintenance-opening phase, revealed that RACK1A likely acts on hook opening via mechanisms involving ARF7 and ARF19 signaling, at the level of auxin-ethylene crosstalk. Collectively, our results imply that RACK1A regulates the abundance of ARF7 and ARF19, particularly in the inner hook side, leading to suppression of the auxin response maximum in this region and consequent re-establishment of equal growth across the hook, resulting in its opening. We have therefore revealed a new role for RACK1A in promoting the opening phase during apical hook development.

RACK1A is a versatile scaffold protein that has already been shown to be involved in multiple plant hormonal signaling pathways, including that of auxin and ethylene (Islas-Flores et al., 2015). Our work suggests that the auxin-ethylene crosstalk signaling pathways in which RACK1A is involved, that positively regulate hook opening, do so by negatively regulating the auxin response maximum found in the inner side of the hook. In contrast, previous studies have reported a positive association between RACK1A and auxin response in the control of lateral root development (Chen et al., 2006; Alshammari et al., 2021). These opposing findings suggest that RACK1A may mediate auxin response in a flexible way, depending on the cell/tissue type and timing of development, implying a role in spatio-temporal regulation of the ARF machinery. Interestingly, although our work suggests that RACK1A enhances ARF7 and ARF19 protein abundance in the apical hook, RACK1A has been proposed as a down-regulator of auxin signaling during salinity acclimation (Denver and Ullah, 2019). Although our results suggest that functional EIN2 is required for RACK1A to induce hook opening, a previous report showed that EIN2 does not interact physiologically with RACK1 proteins in yeast (Wang et al., 2019). Our study also suggests that the transcription factor EIN3 may be required for the mechanism through which RACK1A regulates hook opening, and recent work has suggested that RACK1A binds to EIN3 to enhance its DNA binding activity in *Arabidopsis* (Masood et al., 2023). HLS1 has already been established as a central ethylene-responsive player in the hook formation phase at the level of crosstalk with auxin, specifically acting downstream of EIN3 and EIL1 and upstream of ARF2 (Lehman et al., 1996; Li et al., 2004; An et al., 2012). Our work suggests that HLS1 may play a role in the hook opening process through a different signaling pathway, also possibly involving EIN3 and EIL1, but additionally EIN2, RACK1A, ARF7 and ARF19. The apparent flexibility of these various signaling components, that seem to interact in different ways depending on the developmental context, suggest an effective, adaptable means by which plants ensure plasticity, with appropriate developmental responses to complex signaling networks. Considering that RACK1A is a scaffold protein, it may integrate unknown interactors to modulate the EIN2-mediated ethylene signaling cascade during its regulation of the auxin response in the apical hook. Identification of such missing components will facilitate a better understanding of the mechanisms mediating differential growth during apical hook development.

## Materials and Methods

### Plant materials and growth conditions

All *Arabidopsis thaliana* mutants and transgenic lines were in Columbia-0 (Col-0) ecotype, except *shy2-2* and *gai-1*, which were in Landsberg *erecta* (Ler) ecotype. *Arabidopsis* lines *axr1-30* (Hotton et al., 2011), *DR5::GUS* (Ulmasov et al., 1997), *tir1-1afb2-3* (Parry et al., 2009), *axr2-1* (Nagpal et al., 2000), *shy2-2* (Tian and Reed, 1999), *arf2-8* and *arf7-1arf19-1* (Okushima et al., 2005), *ARF19::ARF19-Venus* and *ARF7::ARF7-Venus* (Powers et al., 2019), *ein2-t* (Fischer et al., 2006), *ein3-1eil1-1* (An et al., 2012), *etr1-1* (Bleecker et al., 1988), *hls1-1* (Lehman et al., 1996), *EBS::GUS* (Stepanova et al., 2007), *gai-1* (Koornneef et al., 1985), *wag2-1* (Santner and Watson, 2006), *ahk2-2ahk3-3* (Higuchi et al., 2004), *rack1a-3* and *amiR-rack1-es1* (Cheng et al., 2015) and *DR5::n3GFP* (Liao et al., 2015) were described previously. For experiments in which multiple genotype combinations of *axr1-30*, *rack1a-3* and *ein2-t* were used (Fig. 5B to E and *SI Appendix*, Fig. S9), they were assembled by genetic crosses and were also combined with *DR5::n3GFP*, as were all control lines including WT and single mutants. The homozygous lines were screened using fluorescence and PCR genotyping, which was performed as previously described (Edwards et al., 1991) using the following primers. For *axr1-30* (SAIL_904_E06) T-DNA mutant: wild-type allele, 5’-CAAACCTGTTTTCTCTTCCAGG-3’ and 5’-ATCTGTCCGCAGCTCTAAGAAG-3’; T-DNA allele, 5’-TCTGAATTTCATAA CCAATCTCGATACAC-3’ and 5’-ATCTGTCCGCAGCTCTAAGAAG-3’. For *rack1a-3* (CS862351) T-DNA mutant: wild-type allele, 5’-GGCATCTCCAGACACCGAAA-3’ and 5’-GCAGAGAGCAACGACAGC-3’; T-DNA allele, 5’-TCTGAATTTCATAACCAATC TCGATACAC-3’ and 5’-GCAGAGAGCAACGACAGC-3’. For *ein2-t* (SALK_086500C) T-DNA mutant: wild-type allele, 5’-AAGTCAAGGACACGGTGAATG-3’ and 5’-GGGTGCATATGGTAACACCAC-3’; T-DNA allele, 5’-GGATTTTGCCGATTTCG GAAC-3’ and 5’-GGGTGCATATGGTAACACCAC-3’.

For most experiments, surface-sterilized seeds were sown on solid growth medium (half-strength (2.2 g L^-1^) Murashige and Skoog (MS) medium (Duchefa), 0.05% morpholinoethanesulfonic acid (Sigma-Aldrich), 1% w/v sucrose, 0.7% w/v plant agar (Duchefa), pH 5.6) supplemented with indicated concentrations of DAPIA, DAPIA metabolites and analogs, estradiol or equal volume of DMSO solvent as the mock treatment, and incubated at 4°C for two days for cold stratification. For ARF7 and ARF19 experiments, plant nutrient medium (Haughn and Somerville, 1996) with 0.5% w/v sucrose and 0.6% w/v agar was used. Assays were performed with etiolated seedlings; plates were incubated in white light at 22°C for 6-12 h to simulate seed germination and then vertically positioned in darkness for 4 d (or 10 d for kinematic analysis, as described below).

### Chemical screen

The chemical screen was conducted as described by Vain et al. (2019). A total number of 4,560 diverse compounds from ChemBridge were dissolved individually in dimethyl sulfoxide (DMSO) as 5 mM stock solutions. In wells of 24-well plates, 300 μL solid medium was supplemented with compounds at 17 μM, with DMSO (mock) controls present in each plate. *Arabidopsis* seedlings of Col-0 and *axr1-30* were grown side by side within each well for 5 d under light conditions (described earlier). In the first round, compounds affecting development in the WT, such as primary/lateral root growth, hypocotyl elongation and gravitropism response, but not or to a lesser extent in *axr1-30* seedlings, were selected. Their effects were confirmed in a second round of screening using fresh compound powders. DAPIA was selected from the reproducible hits as a compound of interest, which rescued the apical hook defect in 4-day-old *axr1-30* etiolated seedlings in a dose-dependent manner, without affecting the Col-0 apical hook phenotype at this time point.

### Chemical preparation

DAPIA was isolated in a screen of ChemBridge molecules (https://www.hit2lead.com/) and has Chembridge ID 5327472. We then synthesized DAPIA and the DAPIA analogs used for SAR analysis (*SI Appendix*, Supporting Text) (Xu and Wolf, 2009; Mevellec et al., 2015; Ninkovic et al., 2010). DAPIA-N (3,5-dimethoxybenzamide) was purchased from TCI America (via Fisher Scientific). All other chemicals, including DAPIA-C (3,5-dimethoxybenzoic acid), were purchased from Sigma-Aldrich. Stock solutions of DAPIA, its metabolites and analogs, and estradiol, were prepared in DMSO.

### Kinematic analysis of apical hook development

For kinematic analysis of apical hook development, seedlings were grown vertically on solid medium plates at 22°C in a dark box illuminated with infrared light from 850 nm LEDs. Seedlings were photographed every hour, in order to catch the precise time of germination, for 10 days using a Canon D50 camera without an infrared filter. Hook angle was then quantified by the angle tool in ImageJ every 4 hours starting 12 hours after germination. Data represent mean ± SE. To analyze the effects of treatments or mutations on the late maintenance-opening phase, the sections of the kinematic curves ranging from the time point at which the maximum hook angle was measured in the control to that at which the angle measured in the control first fell below 30% of its maximum, were considered for the statistical comparison of two curves.

### GUS staining

Seedlings were grown on solid medium supplemented with chemicals as indicated for 4 d in darkness and then directly collected for GUS staining, which was performed as previously described (Béziat et al., 2017b). Briefly, seedlings were fixed in 80% acetone at −20°C for 20 min, washed with 0.1M phosphate buffer (Na_2_HPO_4_/NaH_2_PO_4_) three times, and transferred to GUS staining buffer (Na-phosphate buffer, 1 mg mL^-1^ X-Gluc, 0.1% Triton X-100, 10 mM EDTA, 0.5 mM potassium ferrocyanide, and 0.5 mM potassium ferricyanide). Samples were incubated in darkness at 37°C for 9 h, followed by a series of clearing steps. Stained seedlings were then mounted on glass slides in a mixture of chloral hydrate: glycerol: H_2_O (8:3:1) and bright-field images were captured with a Leica DMi8 epifluorescence microscope.

### Confocal laser scanning microscopy and fluorescence quantification

Fluorescence imaging was performed with a Zeiss LSM 880 confocal microscope, using a Plan-Apochromat 10x/0.45 objective, except for ARF7 and ARF19 experiments, for which a Leica DMi8 confocal microscope with 20x objective was used. Images were acquired using identical imaging settings across samples within each experiment. For quantification of *ARF19::ARF19-Venus* fluorescence signal in the inner side of the apical hook, maximum intensity projections of z-stacks were generated and analyzed in ImageJ software (Fiji). A 100×50 µm region was manually selected in the inner hook side, in which the fluorescence signal intensity was measured as the mean gray value. For quantification of *DR5::n3GFP* fluorescence signal ratio across the apical hook, maximum intensity projections of z-stacks were generated in ZEN software (Zeiss) and analyzed in ImageJ software (Fiji). The inner and outer hook sides were manually defined separately as regions of interest (ROIs) using the ImageJ polygon selection tool, starting directly after the bottom of the shoot apical meristem, and extending to an equal distance on the other side of the hook curvature (and using a similar ROI length for open hooks). Two images containing the fluorescent signal from either side were generated. The ImageJ plugin StarDist was used with default settings to measure the nuclear fluorescence signal intensity as the mean gray value in each ROI. The auxin response gradient across the hook was expressed as inner:outer side ratio of fluorescence signal intensity. For the genotypes in which outer and inner sides could not be distinguished due to an open hook, the gradient was expressed as higher:lower fluorescence side ratio.

### Stability of DAPIA in medium and plants

The stability of DAPIA was detected in both growth medium and plants. Col-0 and *axr1-30* seedlings were grown separately on solid medium supplemented with either DMSO or 10 µM DAPIA for 4 d in darkness. The media were collected directly after the solubilization of the chemicals (0 d) and after 4 d in the presence or absence of etiolated seedlings, at the same time point as at which the seedlings were collected. All sample types were flash-frozen in liquid nitrogen and stored at −80°C until extraction. All experiments were done in triplicate. An ACQUITY UPLC I-Class system combined with a Xevo TQ-S triple quadrupole mass spectrometer (Waters, Manchester, UK) was used to quantify DAPIA and its potential metabolites as previously described by Pařízková et al. (2021). Briefly, medium samples were heated in a microwave oven and then diluted 1/100 with methanol. For plant samples, 30 mg of tissue fresh weight was extracted in 100% methanol and purified by liquid-liquid extraction (MeOH:H_2_O:hexane – 1:1:1). Each sample (1 µL) was then injected onto a reverse-phase column (Kinetex C18 100A, 50×2.1 mm, 1.7 μm; Phenomenex) and analyzed by the selected ion recording (SIR) modes for media samples ([M+H]^+^: *m/z* 308, 182 and 183 for DAPIA, DAPIA-N and DAPIA-C, respectively) and multiple reaction monitoring (MRM) modes for plant tissues samples (308 > 165, 182 > 139, 183 > 77 for DAPIA, DAPIA-N and DAPIA-C, respectively). Quantification was performed by external calibration, and the compounds were quantified according to their dilutions and/or estimated recoveries (24.2% for DAPIA, 67.7% for DAPIA-N and 45.3% for DAPIA-C). The limits of detection (signal-to-noise ratio of 1:3) were close to 0.1 pmol or 0.1 fmol for DAPIA and DAPIA-C, and 1 pmol or 10 fmol for DAPIA-N using SIR or MRM modes, respectively. The linear range was at least over 3 orders of magnitude with an *R^2^* coefficient of 0.997 to 0.999. All data were processed by MassLynx V4.1 software (Waters).

### ARF7-Venus and ARF19-Venus detection by immunoblotting

Equal quantities of *ARF7::ARF7-Venus* and *ARF19::ARF19-Venus* 4-day-old etiolated seedlings for each tissue sample were collected under green light and flash frozen in liquid nitrogen. Tissue was homogenized using a 2010 Geno/Grinder (SPEX Sample Prep) at 1500 RPM for 30 sec and then placed at 70°C in 2x NuPage LDS buffer (141 mM Tris, 2% lithium dodecyl sulfide, 0.51 mM EDTA, 10% glycerol, 0.175 mM Phenol Red, 0.22 mM Coomassie blue) and loaded onto a Bolt 8% Bis-tris protein gel (Thermo Scientific). The gel was run at 80 V for 15 min and then 160 V for 1 h until proper separation of bands were obtained. The gel was then transferred to a nitrocellulose membrane (Amersham Protan 0.45 µm NC) using a wet-transfer system. The gel was washed with Ponceau solution (0.1% w/v Ponceau, 5% acetic acid) for 2 min, washed in TBS-T solution (200 mM Tris, 1.5 M NaCl, 0.1% Tween 20) until excess Ponceau solution was removed, and imaged for loading control analysis. The membrane was then blocked in 8% milk in TBS-T buffer for 1 h before incubation in rabbit anti-YFP antibody (Agri-sera) at 1:5000 overnight at 4°C. The membrane was then washed 3 times for 5 min in TBS-T solution before being incubated with anti-rabbit HRP-coupled secondary antibody at 4°C overnight. The signal was then detected using a WesternBright ECL HRP substrate kit (Advasta) according to the manufacturer’s instructions.

### DARTS assay and RACK1A detection by immunoblotting

The DARTS assay for target identification and validation of potential protein interactors of DAPIA were performed as described (Lomenick et al., 2011) with some modifications. For unbiased target identification, *Arabidopsis* 4-day-old etiolated seedlings were used for total protein extraction. All extraction steps were conducted at 4°C. After harvesting in green light, 0.5 g tissues were ground in liquid nitrogen, resuspended in 1 mL total protein extraction buffer (10 mM Phosphate Buffered Saline (PBS) at pH 7.4 (Sigma P5368), 0.5% v/v NP-40 (Nonidet™ P 40 Substitute from Sigma 74385), 2 mM DTT, 1x protease inhibitors (Roche 11836153001) and 1x phosphatase inhibitors (Roche 04906845001) at a 1:2 w/v ratio, and centrifuged to discard the cell debris. After determining the protein concentration with Bradford protein assay reagents (Sigma B6916), the protein extract at 5 mg mL^-1^ was split into two LoBind 1.5 mL tubes and incubated with 100 µM DAPIA or equal volume of DMSO (1% v/v) as mock control for 1 h at room temperature with slow mixing. The compound concentration used was much higher than the biologically relevant dose in order to saturate the protein with ligand and ensure maximal protection from proteolysis (Lomenick et al., 2011). The treated protein extracts were further aliquoted, and each of the aliquots was mixed with pronase (Roche) (1.25 mg mL^-1^ stock solution) at the corresponding dilution to achieve the aimed ratio of total enzyme to total protein substrate: 1:100, 1:300 and 0 (no pronase). After incubation for 30 min at room temperature, the proteolytic digestion was stopped by adding protease inhibitor cocktail (PIC) (Roche) and the tubes were placed on ice immediately. To prepare samples for proteomic analysis, all the following steps were performed on-column using Vivacon 500 10K Spin Columns (Sartorius Stedim VN01H01). The DARTS protein samples were denatured and reduced in denaturing buffer (6 M guanidine, 0.1 M Tris and 5 mM EDTA, pH 8.0) containing 0.3% w/v DTT at 70°C for 1 h, alkylated in alkylation buffer (1.5% w/v iodoacetamide in denaturing buffer) at room temperature for 30 min in the dark, and digested by trypsin (Promega V5280) at 1:100 w:w enzyme-to-sample ratio for each sample in 50 mM ammonium bicarbonate, pH 8.0, at 37°C for 16 h in the dark with gentle rotation. After column centrifugation, the flow-through was collected, containing the tryptic peptides to be analyzed by LC-MS/MS, leading to the identification of RACK1A as a potential target of DAPIA.

For validation of the ligand binding to its potential target, the compound-treated protein extracts were divided into six aliquots of 50 μL, to which different dilutions of pronase were added as indicated and digested for 30min at room temperature. After mixing with 4x Laemmli sample buffer (Bio-Rad 1610747) containing 2-mercaptoethanol and boiling at 70°C for 10 min to stop the proteolytic reaction, the protein samples were loaded onto SDS-PAGE gels and immunoblotting was performed. Membranes were probed with either rabbit anti-RACK1A (1:500) (Chang et al., 2005) or mouse anti-α-Tubulin (1:3000) (Sigma-Aldrich T6074) antibodies. The secondary antibodies were HRP-coupled goat anti-rabbit (Agrisera AS09602) and goat anti-mouse (Santa Cruz sc-2005). Blots were developed with SuperSignal West Dura Extended Duration Substrate (Thermo Scientific 34075) according to the manufacturer’s instructions, or stained with Coomassie blue, and imaged with a Bio-Rad ChemiDoc XRS+ molecular imager.

### LC−MS/MS analysis of tryptic peptides

A 1 μg aliquot of each trypsin digested sample was loaded on a BEH C18 analytical column (75 μm internal diameter × 250 mm, 1.7 μm particles; Waters, MA, USA) and separated using a concave 180 min gradient of 1–40% solvent B (0.1% formic acid in acetonitrile) in solvent A (0.1% aqueous formic acid) at a flow rate of 368 nL min^-1^. The eluate was passed to a nano-electrospray ionization-equipped Synapt G2-Si HDMS mass spectrometer (Waters, MA, USA) operating in a resolution mode. All data were collected using ion-mobility-MS^E^ with dynamic range extension enabled using a scan time of 0.4 s, mass-corrected using Glu-fibrinopeptide B and Leu-enkephalin as reference peptides.

The LC-MS/MS data were processed with Protein Lynx Global Server v.3.0.3 (Waters, MA, USA), and the resulting spectra were searched against the *Arabidopsis* TAIR10 database. The database search settings were: enzyme-specific cleavage with one miscleavage allowed; carbamidomethylated cysteines as fixed modification; oxidized methionine, N-terminal acetylation, and deamidated asparagine and glutamine as variable modifications. A minimum of three fragments were required for peptide detection with a precursor and fragment tolerance of 10 and 25 ppm, respectively, with a false discovery rate <5%.

### Molecular docking

The 3D structure of the DAPIA ligand was generated with Marvin 18.24.0, Chemaxon (https://www.chemaxon.com). The PDB-entry 3DM0 contained the crystal structure of *Arabidopsis* RACK1A as a C-terminal fusion with the maltose binding protein (MBP). A pdb file containing only the RACK1A structure, encompassing residues Leu5 to Ile324 of the protein’s 327 amino acids, was generated with the open-source software PyMOL (version 1.7, Schrödinger, LLC) by removing the MBP sequence and used for computational docking. AutoDockTools version 1.5.6 (Forli et al., 2016) was used for pdbqt-format preparation of the protein and ligand. Docking simulations were executed with AutoDock 4.2 (Morris et al., 2009). A blind docking encompassing the full protein structure was first performed to predict one or more optimized poses for the ligand, followed by local docking focusing on the optimized pose(s) area to find the most favorable pose. The default parameters in AutoDock that are suitable for most drug-sized ligands were used, except ‘ga_pop_size’ and ‘ga_run’ set to 300 and 500, respectively. For blind dockings, the grid-box size was x, y and z = 126 with Grid Point Spacing = 0.453 Å centered at x = 54.959, y = 34.163, and z = 31.695, while x, y = 70 and z = 112 with 0.375 Å centered at x = 54.959, y = 36.284, and z = 31.695 for local dockings. The binding site was predicted by the conformational cluster with the lowest estimated free energy of binding and the highest number of runs. The pose of the ligand with the lowest estimated free energy within the binding site was selected and displayed graphically. UCSF Chimera (Pettersen et al., 2004) (https://www.rbvi.ucsf.edu/chimera) was used for visualization.

### Microscale thermophoresis (MST) analysis

MST experiments were carried out using a Monolith NT.115 (NanoTemper Technologies GmbH). Purified insect cell-expressed RACK1A was fluorescently labeled with Protein Labeling Kit RED-NHS 2nd Generation (NanoTemper Technologies GmbH MO-L011) via amine conjugation. Increasing concentrations of titrant (DAPIA or its inactive analogs) were titrated against constant concentrations (30 nM) of the labelled RACK1A protein in a standard MST buffer (50 mM Tris, pH 7.5, 150 mM NaCl, 10 mM MgCl_2_, 0.05% Tween 20). The small molecules DAPIA and its inactive analogs DAPIA-02 and DAPIA-07 were dissolved in DMSO for a final concentration of 5% v/v when added to an equal volume of target protein solution. MST premium-coated capillaries (Monolith NT.115 MO-K025) were used to load the samples into the MST instrument. The LED power and MST power were set at 60% and 40%, respectively, for all the thermophoresis measurements. Before the quantitative binding affinity assays, a binding check procedure was performed to analyze whether the experiment set-up was correct and a binding event could be detected (yes/no answer). MST experiments were performed at room temperature and standard deviation was calculated from four independent replicates. Data were analyzed using the MO.Affinity Analysis 3 software provided together with the instrument by the company and dissociation constant (K_d_) was calculated by the K_d_ fitting function of the software.

### Statistical analysis

For statistical analysis of differences among multiple sample groups, one-way ANOVA and Tukey’s or Duncan multiple comparison tests were performed and different letters in the figures represent significant differences at *P* < 0.05. For statistical analysis of differences between two sample groups, the Student’s T-test and Wilcoxon rank sum test were performed for parametric and non-parametric datasets, respectively. The Compare Groups of Growth Curves (CGGC) method, which performs permutation tests to compare curves of measurements over time (Elso et al., 2004), was used to statistically analyze differences between apical hook angle kinematic curves. This method calculates all pairwise comparisons with T-tests at each time point and averages them to obtain the permutation *P*-value (Elso et al., 2004). Statistical differences are represented as follows: \*\*\**P* < 0.001, \*\**P* < 0.01, \*\*\**P* < 0.05, ns - no statistical difference. Error bars reflect either standard deviation (SD) or standard error of the mean (SEM), as indicated in the figure legends. Biological replicates were performed on different days. Box plots, line graphs and bar graphs were drawn using Microsoft Excel or GraphPad Prism 8.0.2.

## Acknowledgments

We acknowledge the UPSC Microscopy Facility (Umeå, Sweden). The RACK1A antibody was a gift from Julia Bailey-Serres. We thank Jose M. Alonso and Anna Stepanova for critical reading of the manuscript. This work was financially supported by the Kempe Foundation (Q.M., S.L., D.K.B.), the Tryggers Foundation (Q.M.), the Swedish Research Council (Vetenskapsrådet) (S.Ro., S.M.D.), the Knut and Alice Wallenberg Foundation (S.L., S.Ra.), VINNOVA (Verket för Innovationssystem) (S.L., S.Ra.), the Internal Grant Agency of Palacký University (B.P., O.N.) and the National Institutes of Health (E.G.W., L.C.S).

