## Supplementary Information Appendix for "Identification of RACK1A as a component of the auxin-ethylene crosstalk regulating apical hook development in *Arabidopsis thaliana*"

**This PDF file includes:**

- Supporting text
- Figures S1 to S9
- Legend for Dataset S1

**Other supporting materials for this manuscript include the following:**

- Dataset S1

**Supporting Information Text**

**Synthetic procedures for DAPIA, DAPIA-01, DAPIA-02, DAPIA-03, DAPIA-05, DAPIA-06, DAPIA-07, DAPIA-08, DAPIA-09 and DAPIA-10.**

DAPIA-C (CAS: 1132-21-4) and DAPIA-N (CAS: 17213-58-0) were purchased.

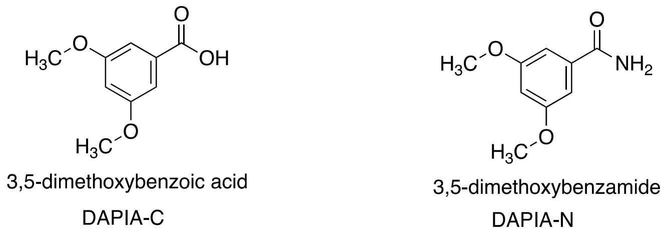


DAPIA-C was used as a precursor to synthesize DAPIA-01 and DAPIA-02. DAPIA-02 was used to synthesize DAPIA-03 and DAPIA-09. DAPIA-05 was used to synthesize DAPIA-08 and DAPIA-08 was used to synthesize DAPIA-10.

**General Experimental procedure for DAPIA, DAPIA-01, DAPIA-02, DAPIA-05, DAPIA-06 and DAPIA-07.** A two neck, round-bottomed flask fitted with reflux condenser and equipped with a magnetic stirring bar charged substituted carboxylic acid (100-500 mg scale) and $CH_2Cl_2$ (20-100 mL). The flask was closed with a rubber septum and placed under nitrogen by piercing the septum with a needle. Oxalyl chloride (2 equiv.) was added dropwise *via* a syringe through the septum followed by the addition of a few drops of DMF. The resulting mixture was refluxed for 6 h (monitored by TLC) and then concentrated on a rotary evaporator. To the resulting crude acid chloride, THF (10-50 mL) was added. To this solution, diisopropylethylamine (DIPEA) (2 equiv.) was added and the mixture was cooled in an ice bath to 0°C and stirred at the same temperature for 30 min. To this mixture, substituted amine (1.5 equiv.) was added and the reaction mixture was stirred at room temperature for 12 h. The mixture was treated with 2 N HCl (adjusted pH to 7), transferred to a separatory funnel and extracted with ethyl acetate. The organic layers were washed with brine and dried over sodium sulfate, concentrated under reduced pressure to give crude material, which was purified by crystallization (ethyl acetate and hexane) or trituration with hexane to afford the desired product (48 to 80%).

**Synthesis of DAPIA:**

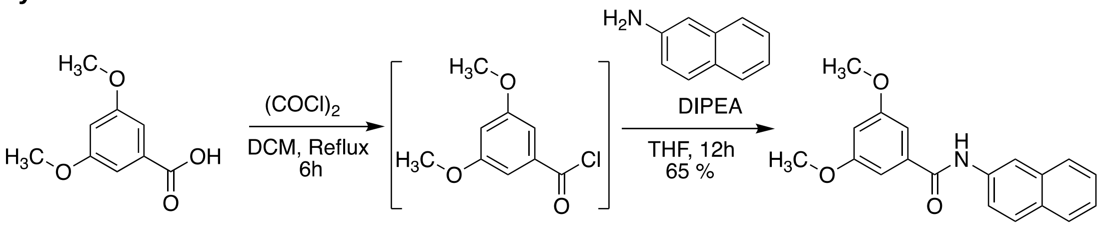


**Experimental procedure for DAPIA.** A two neck, 25 mL round-bottomed flask equipped with a magnetic stirring bar charged 3,5-dimethoxybenzoic acid (50 mg, 0.274 mmol) and $CH_2Cl_2$ (10 mL). The flask was closed with a rubber septum and placed under nitrogen by piercing the septum with a needle. Oxalyl chloride (0.047 mL, 0.069 g, 0.549 mmol) was added dropwise *via* a syringe through the septum followed by the addition of 1 drop of DMF. The resulting mixture was refluxed for 6 h (monitored by TLC) and then concentrated on a rotary evaporator. To the resulting crude acid chloride, THF (10 mL) was added. To this solution, diisopropylethylamine (DIPEA) (0.095 mL, 0.071 g, 0.549 mmol) was added and the mixture was cooled in an ice bath to 0°C and stirred at the same temperature for 30 min. To this mixture, anthracene-2-amine (0.05 g, 0.274 mmol) was added, and the reaction mixture was stirred at room temperature for 12 h. The mixture was treated with 2 N HCl (adjusted pH to 7), transferred to a 50 mL separatory funnel and extracted with ethyl acetate (25 mL x 3). The organic layers were washed with brine (25 mL), dried over sodium sulfate, concentrated under reduced pressure and triturated with hexane to afford DAPIA as an off-white solid (55 mg, 65%).

**Synthesis of DAPIA-01:**

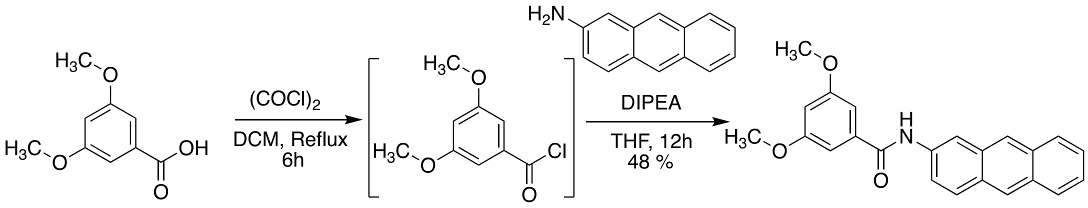


**Experimental procedure for DAPIA-01.** A two neck, 50 mL round-bottomed flask equipped with a magnetic stirring bar charged 3,5-dimethoxybenzoic acid (100 mg, 0.548 mmol) and $CH_2Cl_2$ (20 mL). The flask was closed with a rubber septum and placed under nitrogen by piercing the septum with a needle. Oxalyl chloride (0.094 mL, 0.139 g, 1.09 mmol) was added dropwise *via* a syringe through the septum followed by the addition of 1 drop of DMF. The resulting mixture was refluxed for 6 h (monitored by TLC) and then concentrated on a rotary evaporator. To the resulting crude acid chloride, THF (20 mL) was added. To this solution, diisopropylethylamine (DIPEA) (0.191 mL, 0.142 g, 1.09 mmol) was added and the mixture was cooled in an ice bath to 0°C and stirred at the same temperature for 30 min. To this mixture, anthracene-2-amine (0.159 g, 0.823 mmol) was added, and the reaction mixture was stirred at room temperature for 12 h. The mixture was treated with 2 N HCl (adjusted pH to 7), transferred to a 100 mL separatory funnel and extracted with ethyl acetate (50 mL x 3). The organic layers were washed with brine (100 mL), dried over sodium sulfate, concentrated under reduced pressure, and triturated with hexane to afford DAPIA-01 as an off-white solid (95 mg, 48%).

**Synthesis of DAPIA-02:**

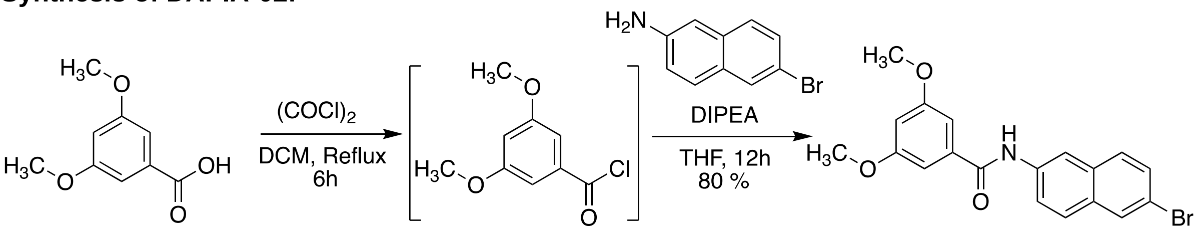


**Experimental procedure for DAPIA-02.** A two neck, 250 mL round-bottomed flask equipped with a magnetic stirring bar charged 3,5-dimethoxybenzoic acid (500 mg, 2.74 mmol) and $CH_2Cl_2$ (100 mL). The flask was closed with a rubber septum and placed under nitrogen by piercing the septum with a needle. Oxalyl chloride (0.47 mL, 0.696 g, 5.48 mmol) was added dropwise *via* a syringe through the septum followed by the addition of 2 drops of DMF. The resulting mixture was refluxed for 6 h (monitored by TLC) and then concentrated on a rotary evaporator. THF was added to the resulting crude acid chloride. To this solution, diisopropylethylamine (DIPEA) (0.949 mL, 0.709 g, 5.48 mmol) was added and the mixture was cooled in an ice bath to 0°C and stirred at the same temperature for 30 min. To this mixture, 6-bromonaphthalen-2-amine (0.914 g, 4.11 mmol) was added and the reaction mixture was stirred at room temperature for 12 h. The mixture was treated with 2 N HCl (adjusted pH to 7), transferred to a 100 mL separatory funnel and extracted with ethyl acetate (50 mL x 3). The organic layers were washed with brine (100 mL) and dried over sodium sulfate, concentrated under reduced pressure, and triturated with hexane to afford DAPIA-02 as an off-white solid (0.85 g, 80%).

**Synthesis of DAPIA-03:**

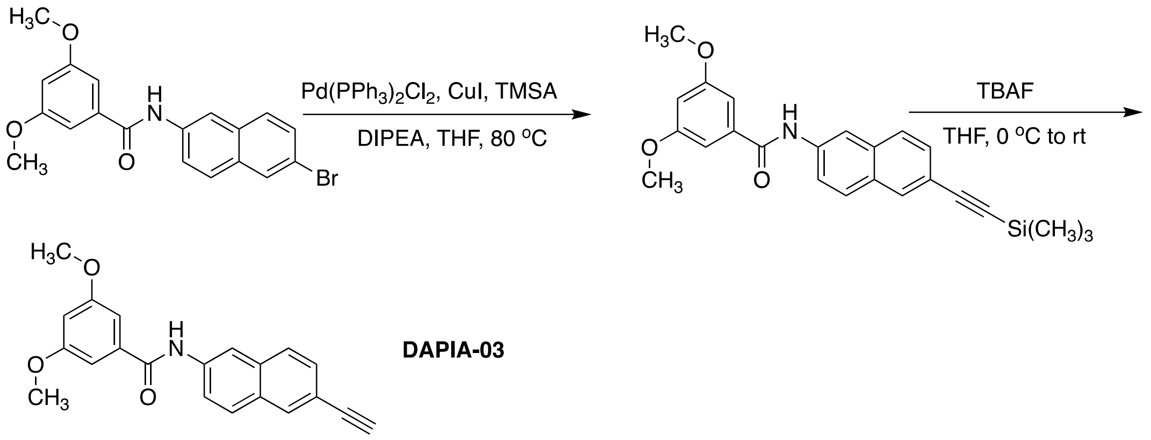

**DAPIA-03**

**Experimental procedure for DAPIA-03.** A mixture of *N*-(6-bromonaphthalen-2-yl)-3,5-dimethoxybenzamide (100 mg, 0.25 mmol), $Pd(PPh_3)Cl_2$ (18.17 mg, 0.025 mmol), CuI (3.94 mg, 0.02 mmol), and diisopropylethylamine (DIPEA) (0.103 g, 0.134 mL, 0.776 mmol) in THF (20 mL) was stirred at 25°C for 30 min under nitrogen. Trimethylsilyl acetylene (TMSA) (0.107 mL, 76.2 mg, 0.776 mmol) was added slowly to the mixture with stirring. The resulting mixture was then stirred at 25°C for 12 h, diluted with water (50 mL), and extracted with ethyl acetate (100 mL x 3). The organic layers were collected, combined, dried over anhydrous $Na_2SO_4$, and concentrated under reduced pressure. The residue was then purified by column chromatography using hexane-ethyl acetate to give the desired product as a low melting solid (80 mg, 76% yield). Sonogashira coupling product was used for the next step for TMS-deprotection. The **coupled product** (50 mg, 0.123 mmol) was taken in dry **THF** (10 mL) and cooled in an ice bath. TBAF (1M in THF, 0.148 mL, 0.148 mmol) was added and the reaction mixture was allowed to warm to RT and stirred for 12 h. The mixture was concentrated *in vacuo*, diluted with ethyl acetate and washed with sat aq. $NH_4Cl$. The organic layer was concentrated to afford crude material, which was purified by column chromatography using ethyl acetate:hexane to afford an off-white solid (20 mg, 48%).

**Synthesis of DAPIA-05:**

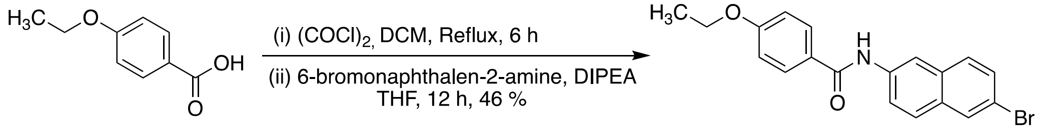

**Experimental procedure for DAPIA-05.** A two neck, 100 mL round-bottomed flask equipped with a magnetic stirring bar charged 4-ethoxybenzoic acid (200 mg, 1.2 mmol) and $CH_2Cl_2$ (30 mL). The flask was closed with a rubber septum and placed under nitrogen by piercing the septum with a needle. Oxalyl chloride (0.206 mL, 305.5 g, 2.4 mmol) was added *via* a syringe through the septum followed by the addition of 2 drops of DMF. The resulting mixture was refluxed for 6 h (monitored by TLC) and then concentrated on a rotary evaporator. To the resulting crude yellow acid chloride, $CH_2Cl_2$ (10 mL) was added. To this solution, diisopropylethylamine (DIPEA) (0.416 mL, 0.311 g, 2.4 mmol) was added and the mixture was cooled in an ice bath to 0°C and stirred at the same temperature for 30 min. To this mixture, 6-bromonaphthalen-2-amine (0.400 g, 1.8 mmol) was added, and the reaction mixture was stirred at room temperature for 12 h. The mixture was treated with 2 $NH_4Cl$ (adjusted pH to 7), transferred to a 100 mL separatory funnel and extracted with ethyl acetate (50 mL x 3). The organic layers were washed with brine (30 mL) and then concentrated under reduced pressure and triturated to afford DAPIA-05 as a white solid (0.205 g, 46%).

**Synthesis of DAPIA-06:**

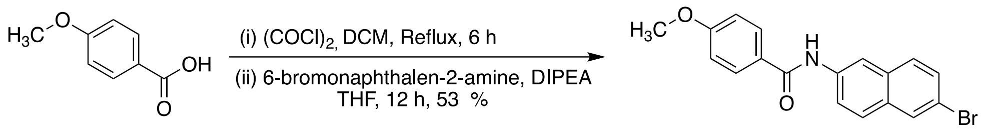


DAPIA-06

**Experimental procedure for DAPIA-06.** A two neck, 50 mL round-bottomed flask equipped with a magnetic stirring bar charged 4-methoxybenzoic acid (100 mg, 0.678 mmol) and $CH_2Cl_2$ (20 mL). The flask was closed with a rubber septum and placed under nitrogen by piercing the septum with a needle. Oxalyl chloride (0.113 mL, 0.167 g, 1.31 mmol) was added *via* a syringe through the septum followed by the addition of 1 drop of DMF. The resulting mixture was refluxed for 6 h (monitored by TLC) and then concentrated on a rotary evaporator. To the resulting crude yellow acid chloride, $CH_2Cl_2$ (10 mL) was added. To this solution, diisopropylethylamine (DIPEA) (0.228 mL, 0.17 g, 1.31 mmol) was added and the mixture was cooled in an ice bath to 0°C and stirred at the same temperature for 30 min. To this mixture, 6-bromonaphthalen-2-amine (0.218 g, 0.986 mmol) was added and the reaction mixture was stirred at room temperature for 12 h. The mixture was treated with 2 N HCl (adjusted pH to 7), transferred to a 100 mL separatory funnel and extracted with ethyl acetate (50 mL x 3). The organic layers were washed with brine (30 mL) and then concentrated under reduced pressure and triturated to afford DAPIA-06 as a white solid (0.125 g, 53%).

**Synthesis of DAPIA-07:**

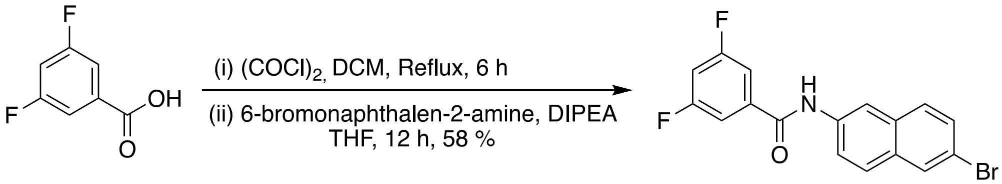


DAPIA-07

**Experimental procedure for DAPIA-07.** A two neck, 50 mL round-bottomed flask equipped with a magnetic stirring bar charged 4-ethoxybenzoic acid (100 mg, 0.632 mmol) and $CH_2Cl_2$ (20 mL). The flask was closed with a rubber septum and placed under nitrogen by piercing the septum with a needle. Oxalyl chloride (0.108 mL, 0.161 g, 1.26 mmol) was added *via* a syringe through the septum followed by the addition of 1 drop of DMF. The resulting mixture was refluxed for 6 h (monitored by TLC) and then concentrated on a rotary evaporator. To the resulting crude acid chloride, $CH_2Cl_2$ (10 mL) was added. To this solution, diisopropylethylamine (DIPEA) (0.219 mL, 0.163 g, 1.26 mmol) was added and the mixture was cooled in an ice bath to 0°C and stirred at the same temperature for 30 min. To this mixture, 6-bromonaphthalen-2-amine (0.211 g, 0.948 mmol) was added, and the reaction mixture was stirred at room temperature for 12 h. The mixture was treated with 2 N HCl (adjusted pH to 7), transferred to a 100 mL separatory funnel and extracted with ethyl acetate (50 mL x 3). The organic layers were washed with brine (30 mL) and then concentrated under reduced pressure and triturated to afford DAPIA-07 as a white solid (0.134 g, 58%).

**Synthesis of DAPIA-08:**

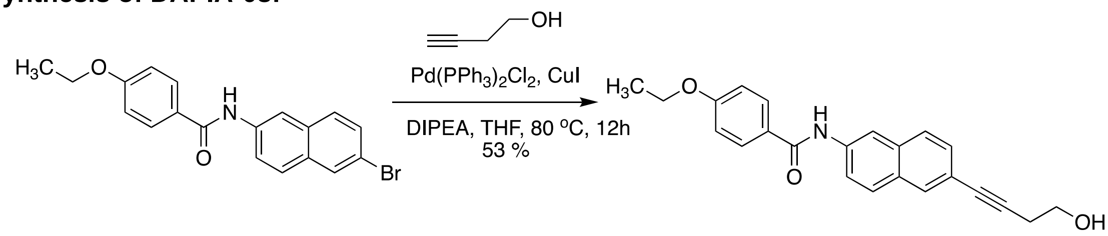

**Experimental procedure for DAPIA-08.** A mixture of *N*-(6-bromonaphthalen-2-yl)-4-ethoxybenzamide (0.2 g, 0.54 mmol), Pd(PPh$_3$)Cl$_2$ (0.038 g, 0.054 mmol), CuI (0.0083 mg, 0.043 mmol) and diisopropyl ethylamine (DIPEA) (0.28 g, 0.209 mL, 1.62 mmol) in THF (8 mL) was stirred at 25°C for 30 min under nitrogen. 3-Butyn-ol (0.122 mL, 113.9 mg, 1.62 mmol) was added slowly to the mixture with stirring. The resulting mixture was then heated at 80°C for 12 h, diluted with water (50 mL), and extracted with ethyl acetate (50 mL x 3). The organic layers were collected, combined, dried over anhydrous $Na_2SO_4$ and concentrated under reduced pressure. The residue was then purified by column chromatography using hexane-ethyl acetate to give DAPIA-08 (98 mg, 53%).

**Synthesis of DAPIA-09:**

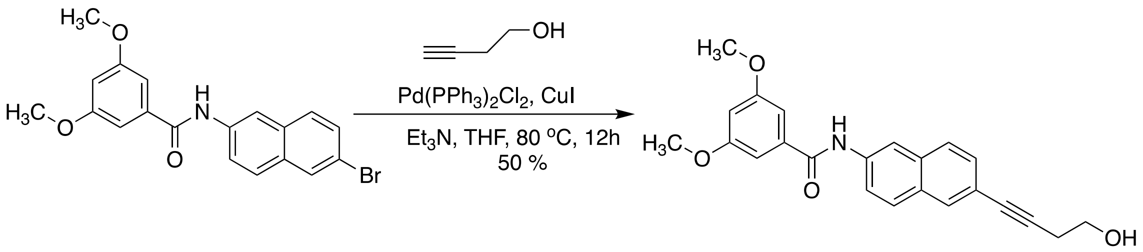


**Experimental procedure for DAPIA-09.** A mixture of *N*-(6-bromonaphthalen-2-yl)-3,5-dimethoxybenzamide (0.1 g, 0.258 mmol), Pd(PPh$_3$)Cl$_2$ (0.018 mg, 0.0259 mmol), CuI (0.0039 g, 0.0207 mmol) and diisopropylethylamine (DIPEA) (0.134 mL, 0.1 g, 0.776 mmol) in THF (8 mL) was stirred at 25°C for 30 min under nitrogen. 3-Butyn-1-ol (0.058 mL, 0.054 g, 0.778 mmol) was added slowly to the mixture with stirring. The resulting mixture was then heated at 80°C for 12 h, diluted with water (20 mL), and extracted with ethyl acetate (25 mL x 3). The organic layers were collected, combined, dried over anhydrous $Na_2SO_4$ and concentrated under reduced pressure. The residue was then purified by column chromatography using hexane-ethyl acetate to afford DAPIA-09 as an off-white solid (50 mg, 50%).

**Synthesis of DAPIA-10:**

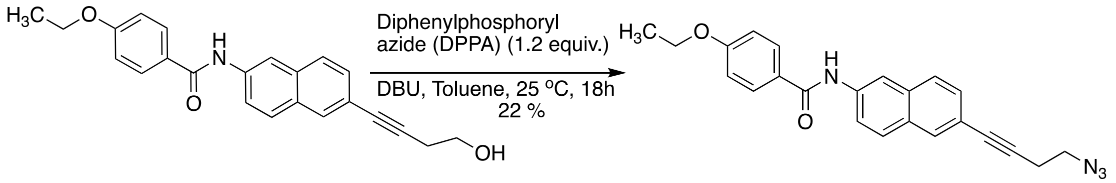


**Experimental procedure for DAPIA-10.** Diphenylphosphoryl azide (0.146 mL, 0.179.9 g, 0.653 mmol) and DBU (0.117 mL, 0.119 g, 0.789 mmol) were added to a suspension of 4-ethoxy-*N*-(6-(4-hydroxybut-1-yn-1-yl)naphthalen-2-yl)benzamide (90 mg, 0.261 mmol) in toluene (50 mL) at 25°C under nitrogen. The resulting mixture was stirred at 25°C for 18 h. The reaction mixture was quenched with sat aq. $NaHCO_3$ (10 mL). The reaction mixture was extracted with EtOAc (150 mL). The combined organics were dried over sodium sulfate and concentrated to a yellow gum. The crude material was purified by column chromatography using ethyl acetate:hexane to provide the product, DAPIA-10, as an off-white solid (22 mg, 22%).

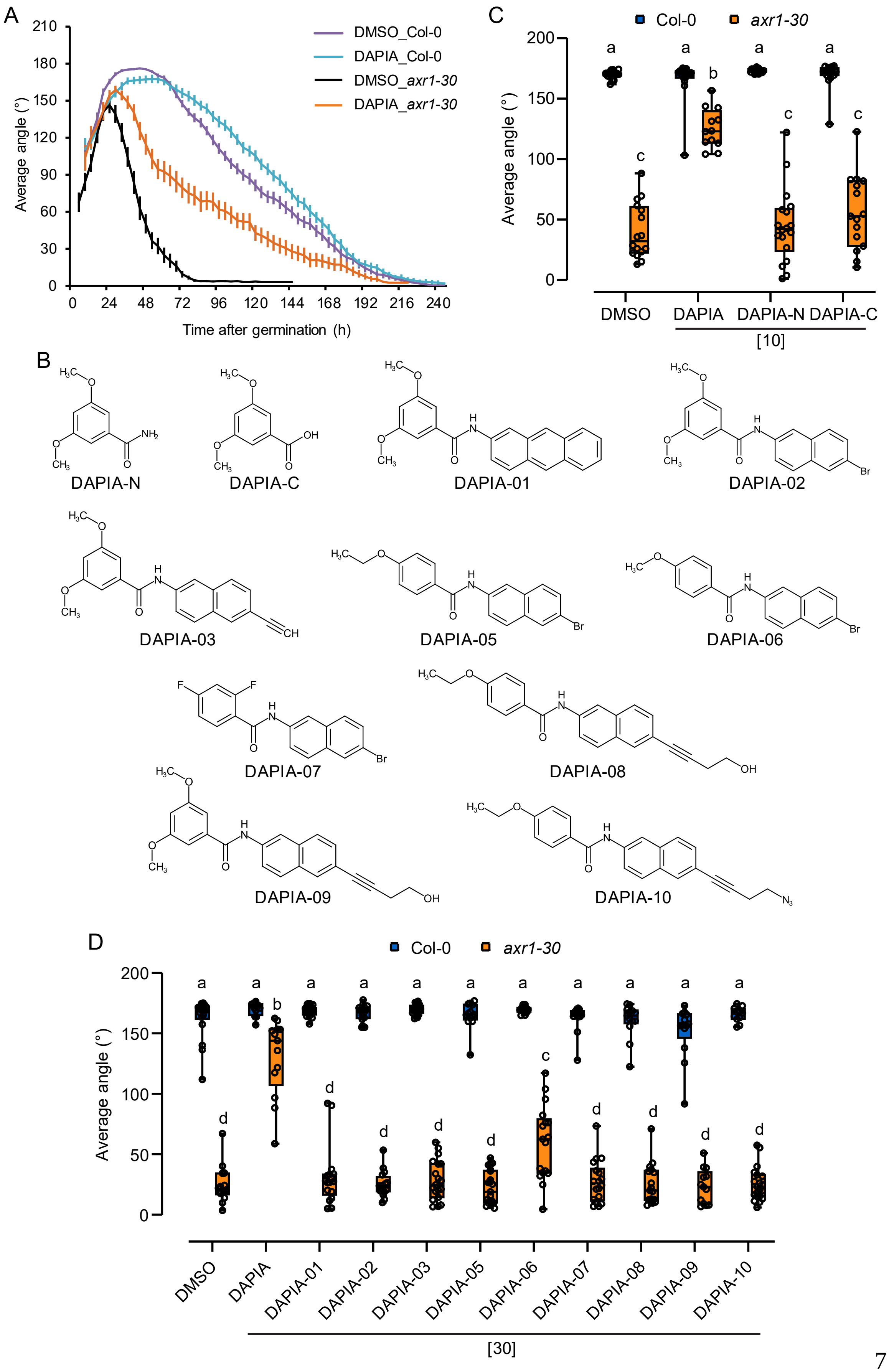
A
Average angle (°)
Time after germination (h)
DMSO_Col-0
DAPIA_Col-0
DMSO_axr1-30
DAPIA_axr1-30
0
24
48
72
96
120
144
168
192
216
240
30
60
90
120
150
180
210
B
DAPIA-N
DAPIA-C
DAPIA-01
DAPIA-02
DAPIA-03
DAPIA-05
DAPIA-06
DAPIA-07
DAPIA-08
DAPIA-09
DAPIA-10
C
Col-0
axr1-30
Average angle (°)
DMSO
DAPIA
DAPIA-N
DAPIA-C
[10]
D
Col-0
axr1-30
Average angle (°)
DMSO
DAPIA
DAPIA-01
DAPIA-02
DAPIA-03
DAPIA-05
DAPIA-06
DAPIA-07
DAPIA-08
DAPIA-09
DAPIA-10
[30]

**Fig. S1.** DAPIA-induced deceleration of apical hook opening and DAPIA structure-activity relationship (SAR). (*A*) Kinematics of apical hook angle in Col-0 and *axr1-30* as measured every 4 hours for 10 days of growth starting from germination (0 h) in darkness on medium supplemented with DMSO (mock) or 10 µM DAPIA. Error bars represent standard error of the mean; n = 23-61 seedlings. (*B*) Chemical structures of potential metabolites and analogs of DAPIA. (*C* and *D*) DAPIA SAR analysis – quantification of average hook angle 4 days after germination and growth of Col-0 and *axr1-30* in darkness on medium supplemented with DMSO (mock), DAPIA and its potential metabolites (*C*) or analogs (*D*). Values in square brackets represent concentrations in µM. Data is represented as box plots and letters indicate significantly different means of n = 11-21 seedlings at $P < 0.05$ (one-way ANOVA, Tukey's multiple comparisons test). Numbers in square brackets represent concentrations in µM.

A

**Medium**

| | DAPIA [µM] | DAPIA-N [µM] | DAPIA-C [µM] |
|---|---|---|---|
| 0d no plants | 9.29 ± 1.54 | nd | nd |
| 4d no plants | 9.90 ± 2.36 | nd | nd |
| 4d Col-0 | 5.17 ± 0.62 | nd | nd |
| 4d *axr1-30* | 4.36 ± 0.09 | nd | nd |

**Plants**

| | DAPIA (pmol/mg FW) | DAPIA-N (pmol/mg FW) | DAPIA -N (%) | DAPIA-C (pmol/mg FW) |
|---|---|---|---|---|
| 4d Col-0 | 472.83 ± 23.12 | 0.021 ± 0.002 | 0.004 | nd |
| 4d *axr1-30* | 476.04 ± 30.90 | 0.022 ±0.002 | 0.005 | nd |

B

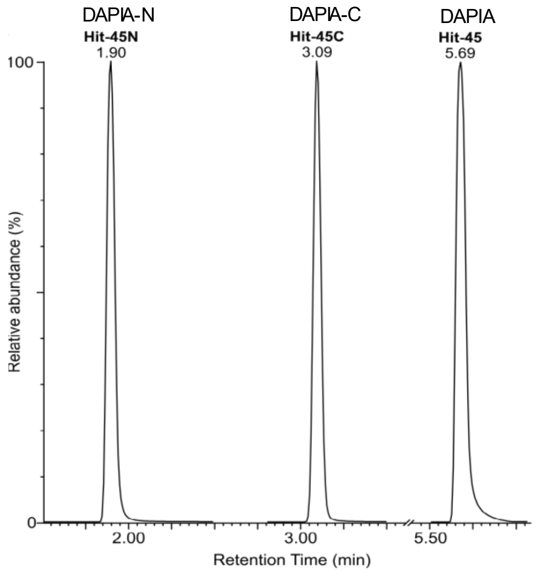


C

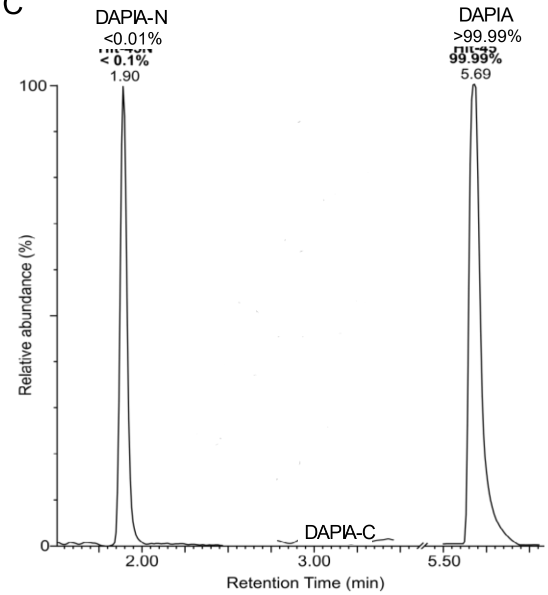


**Fig. S2.** Stability of DAPIA *in vitro* and *in planta*. (*A*) Growth medium was supplemented with 10 µM DAPIA and Col-0 and *axr1-30* seedlings were then germinated and grown on the medium for 4 days in darkness. DAPIA, DAPIA-N and DAPIA-C concentrations were analyzed in the medium directly upon addition of DAPIA (0d) and 4 days later (4d) in the absence or presence of seedlings grown on the medium (Medium) and in the seedlings after the 4 days of growth (Plants). ± values represent standard deviation; nd – not detectable. (*B*) Chromatographic separation of DAPIA (retention time 5.69 min, MRM 308 > 165) and its potential degradation products, DAPIA-N (retention time 1.90 min, MRM 182 > 139) and DAPIA-C (retention time 3.09 min, MRM 183 > 77). (*C*) Representative multi-MRM chromatograms of plant extract. Col-0 and *axr1-30* seedlings were germinated and grown for 4 days in darkness on medium supplemented with 10µM DAPIA. After sample extraction, identification and quantification of analytes were performed using an LC-MS/MS system. DAPIA-N was detected at levels corresponding to less than 0.01% of total DAPIA concentration, while DAPIA-C was not detectable, in the plants. All experiments were done in triplicate.

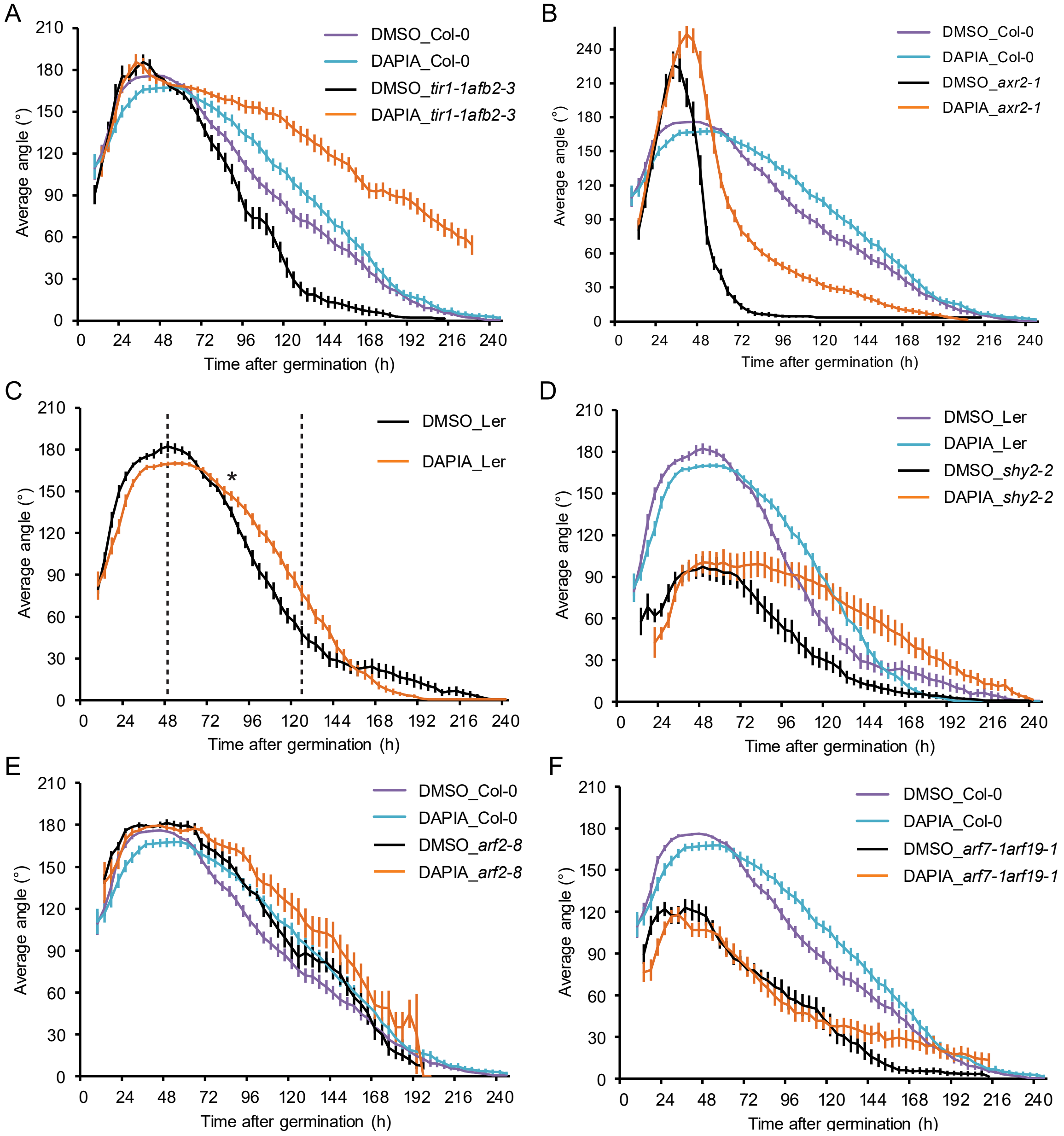


**Fig. S3.** DAPIA requires the auxin signaling components ARF2, ARF7 and ARF19 to decelerate hook opening. (*A* to *F*) Kinematics of apical hook angle in *tir1-1afb2-3* (*A*), *axr2-1* (*B*), *shy2-2* (*D*), *arf2-8* (*E*) and *arf7-1arf19-1* (*F*) together with the relevant WT, as well as in the Ler WT (*C*), as measured every 4 hours for 10 days of growth starting from germination (0 h) in darkness on medium supplemented with DMSO (mock) or 10 µM DAPIA. Error bars represent standard error of the mean; n = 15-61 seedlings. Dashed lines in (*C*) indicate the late maintenance-opening phase of the mock-treated control, for which asterisks indicate significantly different kinematic curves (CGGC method; *$P < 0.05$).

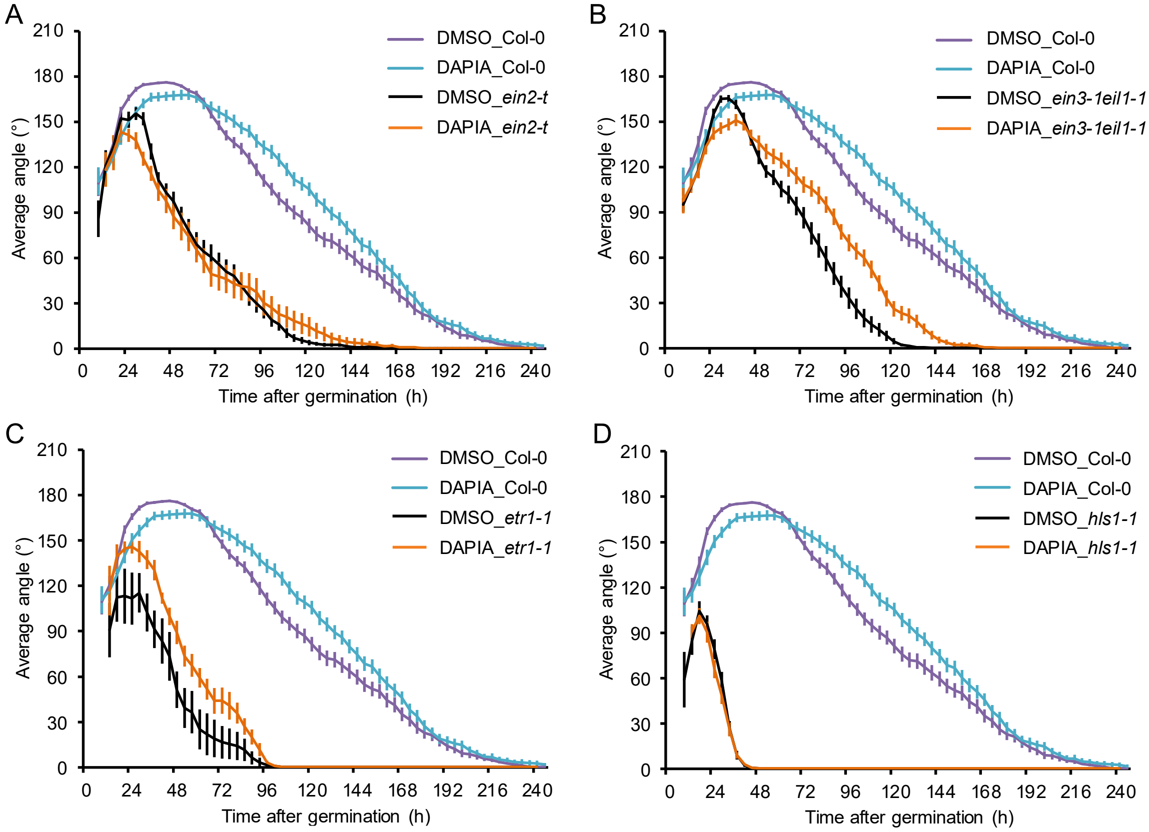


**Fig. S4.** DAPIA requires ethylene signaling components to decelerate hook opening. (*A* to *D*) Kinematics of apical hook angle in *ein2-t* (*A*), *ein3-1eil1-1* (*B*), *etr1-1* (*C*) and *hls1-1* (*D*), together with the Col-0 WT, as measured every 4 hours for 10 days of growth starting from germination (0 h) in darkness on medium supplemented with DMSO (mock) or 10 µM DAPIA. Error bars represent standard error of the mean; n = 7-61 seedlings.

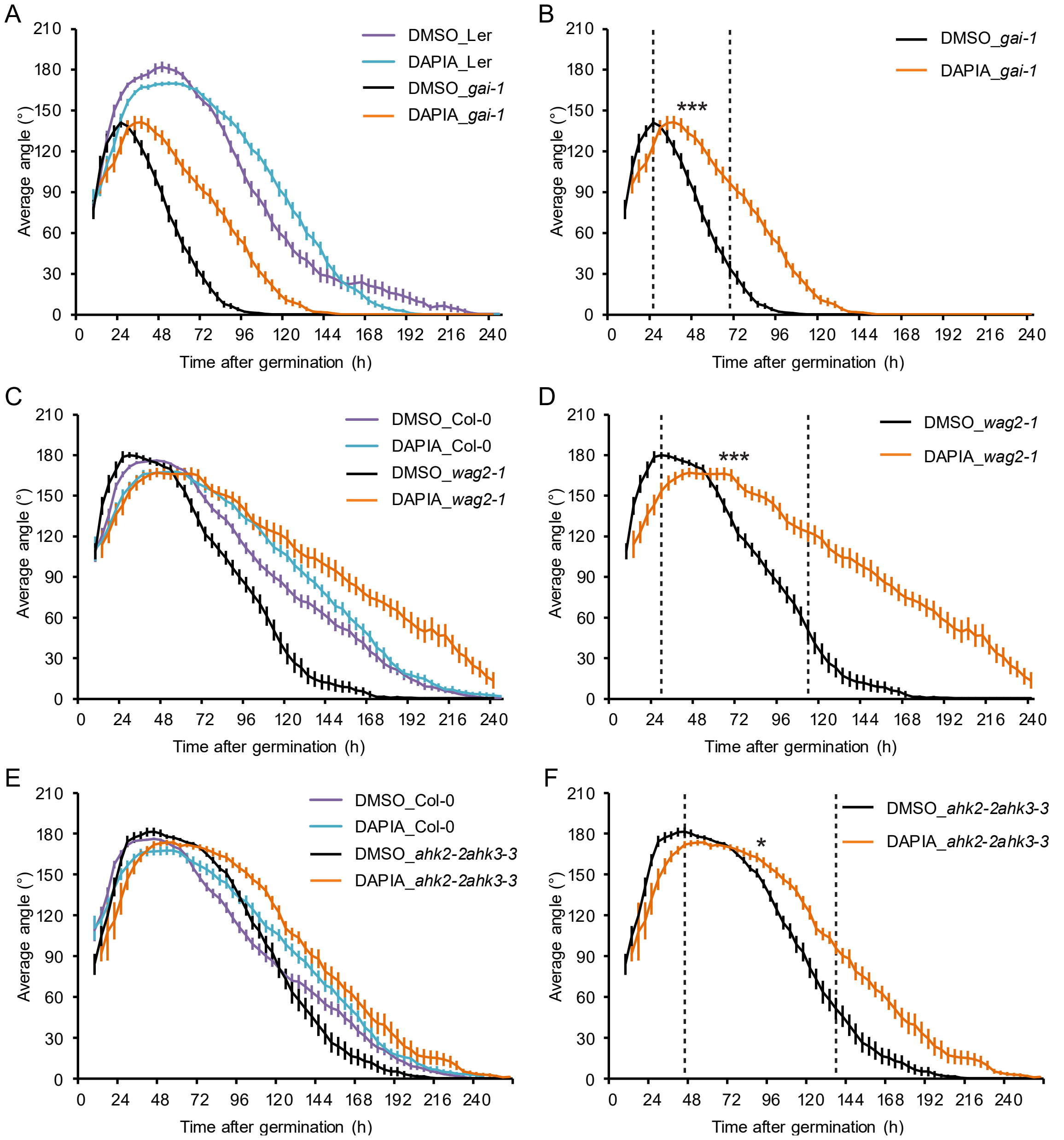


**Fig. S5.** Gibberellin and cytokinin signaling mutants are sensitive to DAPIA. (*A* to *F*) Kinematics of apical hook angle in *gai-1* (*A* and *B*), *wag2-1* (*C* and *D*), and *ahk2-2ahk3-3* (*E* and *F*), together with the relevant WT in (*A*), (*C*) and (*E*), as measured every 4 hours for 10 days of growth starting from germination (0 h) in darkness on medium supplemented with DMSO (mock) or 10 µM DAPIA. Error bars represent standard error of the mean; n = 19-27 seedlings. Dashed lines in (*B*), (*D*) and (*F*) indicate the late maintenance-opening phase of the mock-treated control, for which asterisks indicate significantly different kinematic curves (CGGC method; *$P < 0.05$; ***$P < 0.001$).

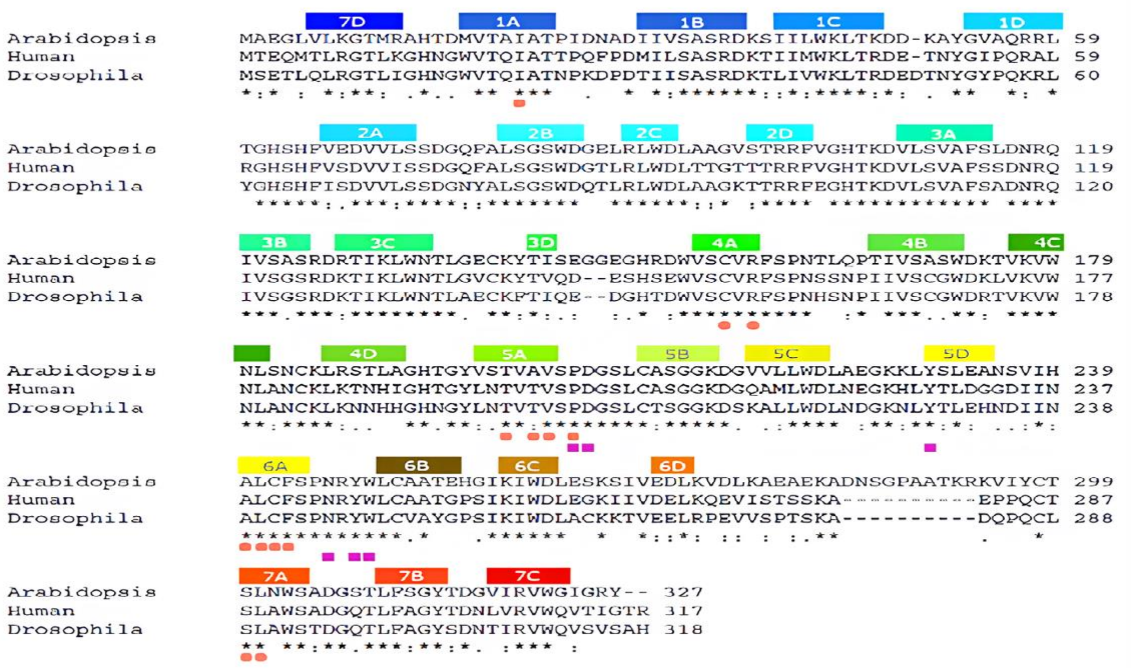


**Fig. S6.** Multiple sequence alignment of *Arabidopsis*, human and *Drosophila* RACK1 in relationship with the 3D structure of *Arabidopsis* RACK1A docked with DAPIA. Protein sequences are available at UniProt (https://www.uniprot.org) with the following accession numbers: *Arabidopsis thaliana* RACK1A: O24456, *Homo sapiens* (human) RACK1: P63244, *Drosophila melanogaster* RACK1: O18640. Sequences are aligned using Clustal Omega. Secondary structural elements of *Arabidopsis* RACK1A (four β-strands in each of the seven blades) are indicated above the sequences, with the same alphanumeric and color codes as used in Fig. 4*B*. Invariant amino acids are denoted by asterisks. The DAPIA-interacting residues are indicated by salmon circles and those constituting the conserved region 2 by purple squares.

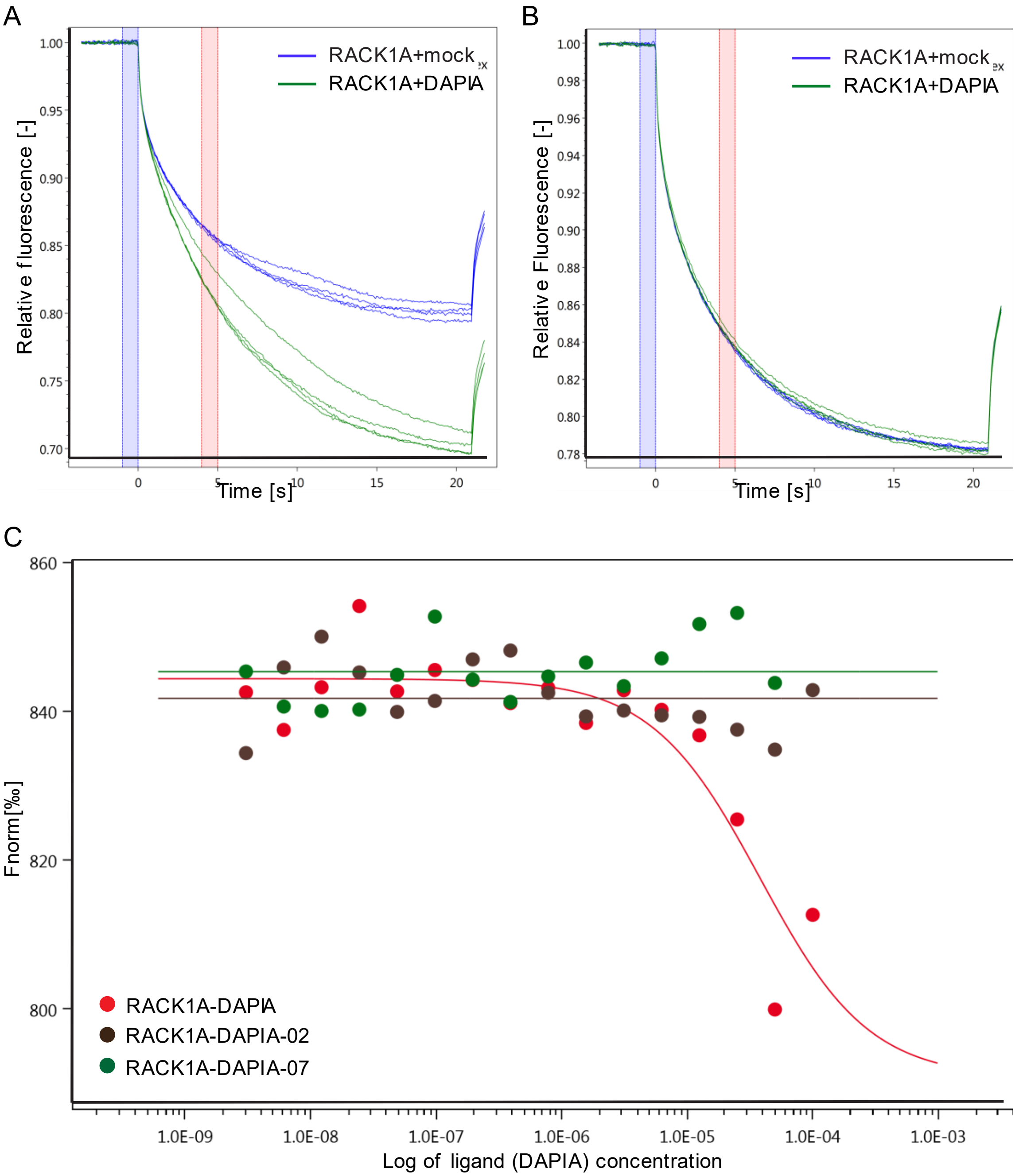


**Fig. S7.** MST analysis validates RACK1A as a target of DAPIA. (*A* and *B*) Qualitative verification of RACK1A and DAPIA binding in the absence (*A*) and presence (*B*) of Tween 20 or PIC. The signal to noise (S/N) value, calculated as the ratio of the response amplitude to the noise of the measurements (standard deviation of the replicates) was 46.9 and 1.4 for DAPIA-treated samples in a and b, respectively. (*C*) Quantification of binding affinity between RACK1A and DAPIA or the inactive DAPIA analogs DAPIA-2 and DAPIA-7.

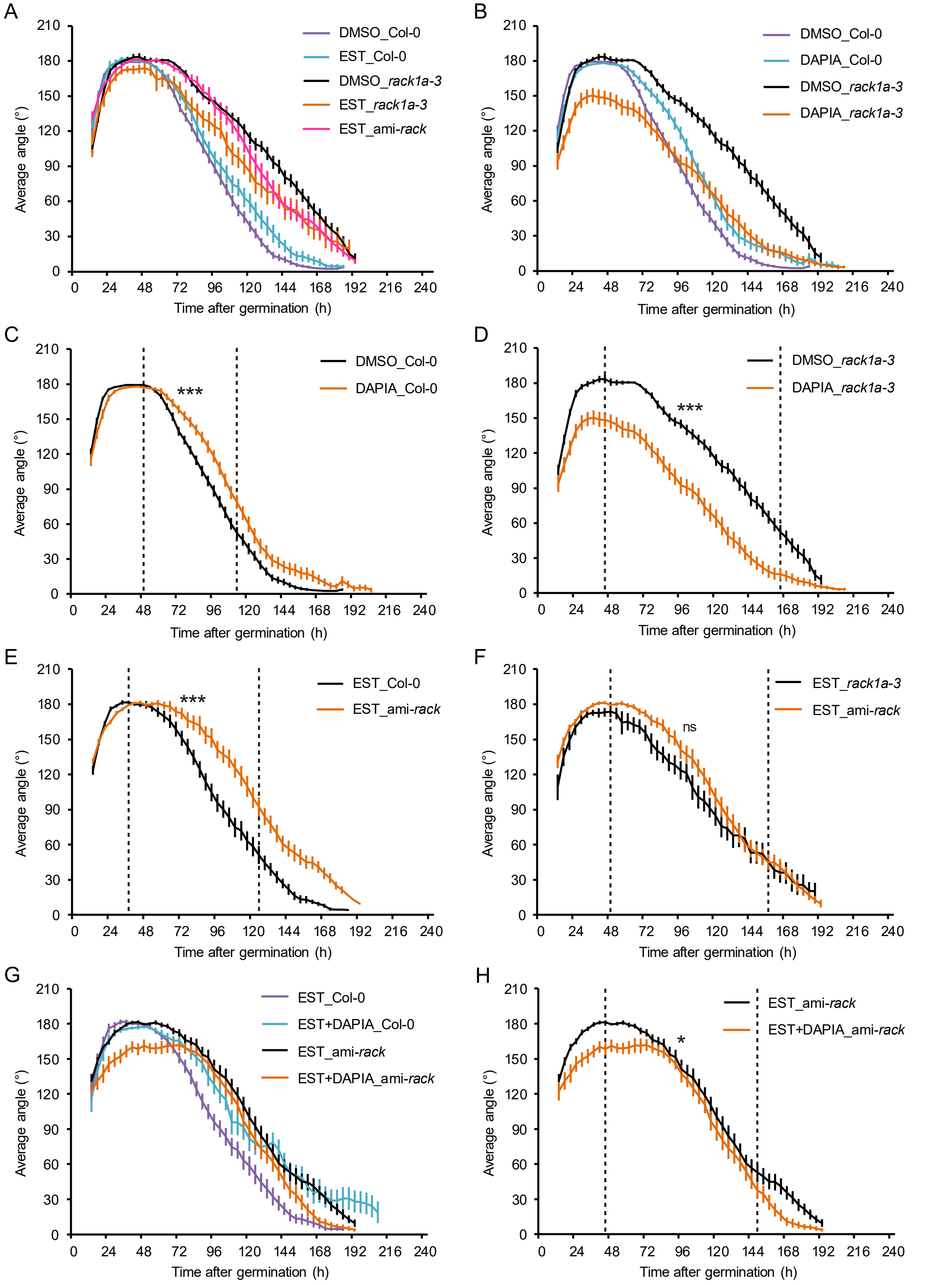


**Fig. S8.** Apical hook development and effects of DAPIA treatment in single and triple mutants of *RACK1*. (*A* to *H*) Kinematics of apical hook angle in Col-0, *rack1a-3* and *amiR-rack1-es1*, as measured every 4 hours for 10 days of growth starting from germination (0 h) in darkness on medium supplemented with DMSO (mock) or 10 µM estradiol, DAPIA, or both. Error bars represent standard error of the mean; n = 14-46 seedlings. Dashed lines in *C* to *F* and *H* indicate the late maintenance-opening phase of the mock-treated control, for which asterisks indicate significantly different kinematic curves (CGGC method; *$P < 0.05$; ***$P < 0.001$).

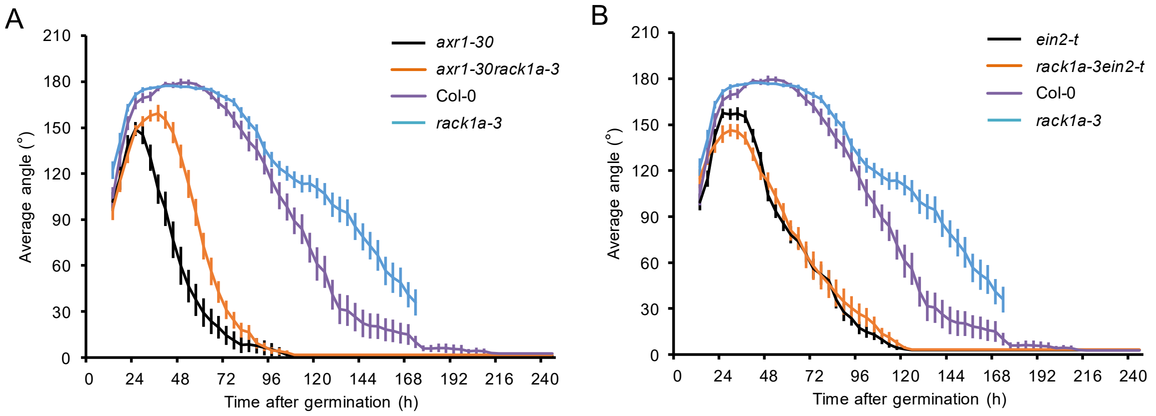


**Fig. S9.** The *rack1a-3* mutation partially rescues the apical hook opening phenotype in *axr1-30* but does not affect hook development in *ein2-t*. (*A* and *B*) Kinematics of apical hook angle in combinations of *axr1-30* with *rack1a-3* (*A*) and *ein2-t* (*B*), together with the single mutants and Col-0 WT, as measured every 4 hours for 10 days of growth starting from germination (0 h) in darkness. Error bars represent standard error of the mean; n = 21-26 seedlings.

**Dataset S1.** Proteins enriched in DARTS assay identified by LC-MS/MS. The spreadsheets Comparison_P100, Comparison_P300 and Comparison_P0 show quantification results for samples subjected to proteolytic digestion by mixing with pronase at enzyme:protein substrate ratios of 1:100, 1:300 or 0 (no pronase), respectively.
